# Transcriptomic-Methylation Integration and Machine-Learning Prioritization of Candidate Prognostic Biomarkers in Triple-Negative Breast Cancer: A Retrospective Computational Study

**DOI:** 10.64898/2026.05.03.722461

**Authors:** Nagendra Mannekunta, Elamathi Natrajan

## Abstract

**Background:** Triple-negative breast cancer is molecularly heterogeneous, complicating prognostic stratification. We aimed to derive a gene panel and clinical-genomic survival nomogram and assess cross-platform performance.

**Methods:** This retrospective computational study analyzed The Cancer Genome Atlas breast cancer data, identifying 197 primary-tumor transcriptomic aliquots from 190 patients. Differential gene-expression and deoxyribonucleic acid (DNA) methylation analyses comparing Dead versus Alive vital status were performed separately. Candidate lists meeting nominal significance (p < 0.05) were intersected by gene symbol to identify a 47-gene discovery signature. Random Forest feature importance, corroborated by LightGBM, prioritized 15 genes. Among 190 patients (163 alive, 27 deceased), standardization and Synthetic Minority Over-sampling Technique resampling yielded 326 observations before an 80%/20% split. Seven classifiers were evaluated; a five-fold sigmoid-calibrated soft-voting ensemble generated the 15-gene risk score. Prognostic performance was assessed using Kaplan–Meier analysis, time-dependent receiver operating characteristic analysis, and a clinical-genomic nomogram. Six immune-infiltration profiles and three immune-checkpoint genes were compared between risk groups. External validation used the GSE58812 microarray cohort (n = 107).

**Results:** Multi-omic integration yielded a 47-gene signature and 15-gene panel. In the 47-gene resampled test partition (n = 66), LightGBM achieved 95.45% accuracy and an area under the receiver operating characteristic curve (AUC) of 0.9899; Support Vector Machine achieved the highest AUC (0.9935). Risk groups differed in overall survival (log-rank p < 0.0001). The risk score remained associated with survival after adjustment for age and advanced stage (hazard ratio 10.67 per standard-deviation increase; 95% confidence interval 6.64–17.15; p < 0.001). Time-dependent AUCs were 0.910, 0.911, and 0.937 at 1, 3, and 5 years. The nomogram estimated 3- and 5-year survival probabilities. Immune-profile (p = 0.38–0.93) and checkpoint-gene differences (p = 0.50–0.74) were not significant. Untouched independent external validation in GSE58812 yielded fixed-model AUCs of 0.4191–0.5500; post-validation XGBoost optimization yielded a gene-only AUC of 0.5533, while an exploratory risk-plus-age score yielded 0.6874.

**Conclusions:** The 15-gene signature showed prognostic association and potentially optimistic internal performance; the nomogram provided a framework for individualized survival estimation. Independent external validation showed limited gene-only transferability; optimized and age-integrated results remain exploratory and require independent assessment.

## Background

Triple-negative breast cancer (TNBC) is a clinically and molecularly heterogeneous subtype of breast cancer, characterized by the absence of estrogen receptor and progesterone receptor expression and human epidermal growth factor receptor 2 (HER2)-negative status [1,2]. This receptor-negative phenotype limits the applicability of endocrine-targeted therapies, while molecular heterogeneity is associated with variable clinical outcomes, highlighting the need for reliable prognostic biomarkers to support patient risk stratification [1,2]. Advances in high-throughput molecular profiling have enabled the integration of transcriptomic and epigenomic data, providing a more comprehensive view of tumor biology than individual molecular data types alone [1,3–5].

Previous TNBC studies have identified correlated regulatory features through multi-omics integration in TNBC cell-line models [4]. In particular, Hashem and Mabrouk [5] integrated transcriptomic and DNA-methylation data to investigate epigenetically regulated metabolic alterations and reported an association between hypermethylated metabolic gene signatures and shorter overall survival. Complementary computational studies have developed prognostic signatures using gene-level mutation features derived from targeted next-generation sequencing [6] and multi-cohort transcriptomic analyses [7]. TNBC accounts for approximately 15–20% of breast cancer cases [7]. Feature-selection approaches have been used to reduce high-dimensional molecular feature sets to focused prognostic signatures [6,7], potentially facilitating the development of assayable biomarker panels. The tumor immune microenvironment is also relevant to TNBC biology and prognosis [2,8]. Immune-related signatures have additionally been associated with response-related characteristics in cancer immunotherapy cohorts [8]. In lung adenocarcinoma, a related study developed an immune-related gene signature for survival prediction and evaluated its association with immune-checkpoint-inhibitor response [9]. Data heterogeneity and model interpretability remain important barriers to the clinical translation of machine-learning approaches in cancer research [10]. Together, these findings provide a rationale for examining selected immune-infiltration and immune-checkpoint features across risk groups.

Although previous TNBC research has demonstrated transcriptomic–epigenomic integration [4,5] and developed computational prognostic signatures [6,7], further evaluation is warranted to determine whether a broader multi-omic discovery signature can be reduced to a focused prognostic panel, whether the resulting molecular risk score can be integrated with clinical variables, and whether its genomic risk signal transfers to an independent cohort measured using a distinct gene-expression platform.

The primary aim of this study was to integrate transcriptomic and epigenomic data from The Cancer Genome Atlas (TCGA) breast cancer cohort (TCGA-BRCA) to derive a multi-omic discovery signature and prioritize a focused candidate prognostic gene panel using Random Forest and LightGBM feature-importance analyses. We further aimed to evaluate the prognostic association and predictive performance of the resulting machine-learning risk score and integrate this molecular score with patient age and American Joint Committee on Cancer (AJCC) tumor stage to construct a clinical-genomic nomogram for estimating individualized 3-year and 5-year overall survival probabilities. Additional objectives were to characterize selected immune-infiltration profiles and immune-checkpoint gene expression across machine-learning-defined risk groups and to evaluate the transferability of the signature-based genomic risk signal through independent cross-platform external validation in the GSE58812 microarray cohort [2].

## Materials and Methods

### 2.1. Cohort Selection and Data Acquisition

This retrospective computational study used public genomic and clinical data to identify prognostic biomarkers in triple-negative breast cancer (TNBC) through transcriptomic–epigenomic integration, machine-learning gene prioritization, and prognostic/predictive evaluation. The internal dataset was obtained from The Cancer Genome Atlas (TCGA) breast cancer cohort (TCGA-BRCA), Data Release 32.0, through the Genomic Data Commons (GDC) Data Portal [1]. The source clinical.tsv comprised 5,546 clinical/treatment records from 1,098 unique patients (cases.submitter_id; one row per recorded therapy regimen), and follow_up.tsv comprised 9,427 longitudinal records from 1,096 patients; the uncollapsed clinical table was retained for repository baseline characteristics (Table 1).

**Table 1.** Baseline characteristics of the uncollapsed source TCGA-BRCA clinical repository.

| Characteristic | Category | Overall |
|---|---|---|
| Source clinical records (unique patients) | n (unique cases.submitter_id) | 5546 (1098) |
| Age at index (years; n = 5545 non-missing) | Mean ± SD | 56.71 ± 12.76 |
| AJCC pathologic tumor stage (n = 4978 non-missing; 568 unreported) | Stage 0 | 12 (0.2%) |
|  | Stage 0is | 3 (0.1%) |
|  | Stage I (I, IA, IB) | 735 (14.8%) |
|  | Stage II (II, IIA, IIB) | 2847 (57.2%) |
|  | Stage III (III, IIIA, IIIB, IIIC) | 1262 (25.4%) |
|  | Stage IV | 77 (1.5%) |
|  | Stage X | 42 (0.8%) |
| Vital status (n = 5545 non-missing; 1 unreported) | Alive | 4697 (84.7%) |
|  | Dead | 848 (15.3%) |
| Overall survival time (days; n = 5540 non-missing) | Mean ± SD | 1340.83 ± 1208.26 |

Eligible primary tumors (sample_type == "Primary Tumor"; shortLetterCode == "TP") were negative for estrogen receptor (ER), progesterone receptor (PR), and human epidermal growth factor receptor 2 (HER2), with Prediction Analysis of Microarray 50 (PAM50) Basal classification [1,2]. Matching yielded 197 primary-tumor RNA sequencing (RNA-seq) aliquots (28-character barcodes), 195 tumor specimens (16-character identifiers), and 190 TNBC patients (12-character identifiers; Supplementary Table 1); five patients had replicate aliquots (3, 3, 2, 2, and 2; four Alive and one Dead). All 197 aliquots (168 associated with Alive and 29 with Dead status) were retained for immune-infiltration and immune-checkpoint analyses (Section 2.4).

For patient-level machine-learning classification, SHapley Additive exPlanations (SHAP) interpretation, and risk-score generation (Section 2.5), patient identifiers (Patient_ID) were extracted from the first 12 characters of the corresponding RNA-seq aliquot barcodes and metadata row identifiers. The expression matrix and corresponding metadata were independently deduplicated by retaining the first-listed occurrence of each Patient_ID and removing subsequent duplicate entries, without averaging replicate expression profiles. This procedure removed seven additional aliquot records attributable to five patients with multiple aliquots, reducing the 197 aliquot-level records to 190 unique patient identifiers. Ensembl gene identifiers were mapped to gene symbols, and the expression matrix was restricted to the predefined 47-gene master biomarker panel, with unmapped and duplicate gene-symbol entries removed. The resulting expression matrix was transposed to represent individual patients as rows and biomarker genes as columns, and patient identifiers were matched against the corresponding clinical metadata. The final patient-level dataset comprised 190 unique patients with complete measurements for all 47 biomarker genes and non-missing Vital_Status (163 Alive [85.8%]; 27 Dead [14.2%]).

Independent external validation used the GSE58812 oligonucleotide microarray cohort (Affymetrix platform) from the National Center for Biotechnology Information (NCBI) Gene Expression Omnibus (GEO) [2,11], comprising 107 TNBC patients (78 alive [72.9%]; 29 deceased [27.1%]) with one primary-tumor expression profile and clinical, demographic, immunohistochemical, and survival metadata. This cohort was used to assess the cross-platform performance of the 15-gene genomic risk signal relative to the TCGA-BRCA RNA-seq discovery dataset.

### 2.2. Multi-Omic Pre-processing and Differential Analysis

Multi-omic preprocessing and differential analyses were performed in R (v4.3.1). Raw TCGA-BRCA RNA-seq counts (60,660 Ensembl identifiers) from Primary Tumor and Solid Tissue Normal samples were analyzed using DESeq2 (v1.38.3) [12]. The corresponding TCGA sample metadata were used to identify the normal reference group by selecting samples annotated as Solid Tissue Normal (sample type code 11; n = 113). All 113 normal reference sample barcodes were present in the RNA-seq count matrix. TNBC Primary Tumor aliquots (n = 197; Supplementary Table 1) were retained as the tumour cohort. DESeq2::vst generated variance-stabilized, normalized log2-scale expression values for visualization, multi-omic correlation, immune profiling, and machine-learning analyses. Exploratory principal component analysis (PCA) visualized Metastatic, Primary Tumor, and Solid Tissue Normal samples (Supplementary Figure 1). Differential expression compared TNBC Primary Tumor aliquots (n = 197; Supplementary Table 1) with Solid Tissue Normal reference samples using negative binomial generalized linear models with an unadjusted single-factor design (∼ sample_type) and no additional clinical or demographic covariates; positive log2FoldChange indicated higher tumor expression. Supplementary Table 2 reports all 60,660 features, including 56,820 with non-missing Wald statistics and nominal p-values and 30,465 with non-missing padj after independent filtering, with baseMean, log2FoldChange, lfcSE, stat, pvalue, and padj.

DNA methylation microarray profiles (Illumina Infinium HumanMethylation450 [450K] BeadChip) were preprocessed using the matched methylation beta-value matrix, which comprised 485,577 cytosine-phosphate-guanine (CpG) probes across 150 samples. Probes containing one or more missing beta values (n = 143,318) were excluded, retaining 342,259 probes with complete beta-value measurements across all samples. All observed beta values in the original matrix were within the valid range of 0–1. For each remaining probe, variance across samples was calculated using the matrixStats package, and probes were ranked in descending order of variance. The 10,000 probes with the highest variance were retained to generate the final methylation matrix for downstream analyses. The resulting matrix contained 10,000 CpG probes across 150 samples, with complete beta values within the valid range of 0–1 (AveExpr 0.0873–0.9163; mean = 0.4633 ± 0.1702). Differential methylation was analyzed using limma (v3.54.2) linear modeling (lmFit) and empirical Bayes moderation (eBayes) [13], comparing Primary Tumor with Solid Tissue Normal using the same unadjusted single-factor design without additional clinical covariates; positive logFC indicated tumor-associated hypermethylation. Supplementary Table 3 reports all 10,000 probes with probe identifier, logFC, AveExpr, t, P.Value, adj.P.Val, and B.

For volcano-plot visualization (Supplementary Figure 2), features were classified as upregulated (log2FC > 1; nominal p < 0.05; n = 326), downregulated (log2FC < −1; nominal p < 0.05; n = 1,720), or non-significant; these thresholds were used only for visualization. In the tumour-versus-normal comparison, 5,097 transcriptomic features met the nominal significance threshold (p < 0.05), including 1,651 with positive log2FoldChange and 3,446 with negative log2FoldChange; 551 features met the Benjamini–Hochberg adjusted p-value threshold (padj < 0.05). Similarly, 571 methylation probes met the nominal significance threshold (P.Value < 0.05), including 450 hypermethylated and 121 hypomethylated probes. No CpG probe reached Benjamini–Hochberg adj.P.Val < 0.05; the minimum adj.P.Val was 0.8709. These tumour-versus-normal findings are distinct from the vital-status-based candidate-gene selection described in Section 2.3.

### 2.3. Integrative Feature Selection and Functional Enrichment

Differential expression analysis of RNA-seq data was performed using DESeq2 to identify genes associated with patient vital status. Patients were classified into Alive and Dead groups, and differential expression was assessed using the contrast Dead versus Alive. Ensembl gene identifiers were mapped to gene symbols using the ‘org.Hs.eg.db’ annotation package. Genes with nominal p-values below 0.05 were retained as RNA-seq candidates.

DNA methylation differences between the same vital-status groups were assessed using the limma package. A linear model was fitted to the selected methylation probes, followed by empirical Bayes moderation. Differentially methylated probes were identified using nominal P-values below 0.05. Probe identifiers were annotated to gene symbols using the Illumina HumanMethylation450K annotation package (‘IlluminaHumanMethylation450kanno.ilmn12.hg19’), using the ‘UCSC\_RefGene\_Nam’ annotation field. Where multiple gene symbols were listed for a probe, the first annotated symbol was retained.

To generate the 47-gene discovery signature, the RNA-seq and methylation candidate gene lists were intersected by gene symbol; a gene was only retained if it met the exact criteria of demonstrating a nominal significance of p < 0.05 in both the RNA-seq and DNA methylation analyses. Duplicate gene symbols were removed, and the corresponding RNA-seq and methylation statistics were combined into an integrated candidate-gene table. This procedure identified 47 overlapping candidate genes, which were retained as the discovery signature for subsequent analyses: ADARB2, ATG9B, BMP4, BOLL, CA10, CACNA1B, CACNA1E, CPEB1, CRCT1, DNAH9, ECEL1, EN2, EYA4, FMO1, FOXE1, GAD2, GCM2, GDF7, HGC6.3, IGLON5, KCNA6, KRT3, LRRC14, LYPD5, MEIS1, MTFR1, MYO7A, NKAIN3, NKX2-2, NPTX2, NXPH1, PAX1, PCSK2, PHACTR1, PPP2R2C, RADIL, SALL1, SDR16C5, SLC1A4, SLC20A2, SLC35F1, SLC7A3, TDRD10, TFAP2E, TLX2, VSX1, and WWP2.

Nominal significance was defined as p < 0.05 for both data types. Multiple-testing-adjusted p-values were also retained and reported to distinguish nominal associations from findings meeting the false-discovery-rate threshold. The 47-gene discovery signature underwent hypergeometric functional enrichment analysis using Enrichr (2024 Web Release) [14] against Reactome 2024, Kyoto Encyclopedia of Genes and Genomes (KEGG) 2026, and WikiPathways 2024. The top 10 pathways from each database, ranked by nominal enrichment p-value, were visualized as −log10(p-value) in Supplementary Figure 3, with p = 0.05 used as the reference threshold.

### 2.4. Immune Microenvironment and Checkpoint Analysis

Single-sample gene set enrichment analysis (ssGSEA) implemented in the Gene Set Variation Analysis (GSVA) package (v1.46.0) [15] was applied to variance-stabilizing transformation (VST)-normalized TCGA-BRCA TNBC expression data for six canonical immune populations: CD8+ T cells (CD8_T_Cells), B cells (B_Cells), M1 macrophages (Macrophages_M1), M2 macrophages (Macrophages_M2), regulatory T cells (Regulatory_T_Cells), and natural killer cells (NK_Cells). The 197 aliquot-level ssGSEA profiles were matched to patient-level 15-gene machine-learning (ML) Risk Scores from 190 unique patients by truncating the 28-character TCGA aliquot barcodes to 12-character patient identifiers and performing an inner join. This matching retained 197 primary-tumor aliquot-level observations across 190 unique patients, including five patients with replicate aliquots. The median ML Risk Score across these 197 matched aliquot-level observations (0.005983164; reported as 0.005983) was used as the cutoff to classify observations into Low-Risk (n = 99; Risk_Score ≤ median) and High-Risk (n = 98; Risk_Score > median) groups. Two-sided Wilcoxon rank-sum tests (wilcox.test; stat_compare_means) were used to compare ssGSEA scores between the Low-Risk and High-Risk groups for all six immune populations. Nominal (unadjusted) p-values were visualized in Supplementary Figure 6.

Targeted immune-checkpoint expression was evaluated from the same genome-wide VST-transformed matrix for CD274 (programmed death-ligand 1 [PD-L1]), PDCD1 (programmed cell death protein 1 [PD-1]), and CTLA4 using version-agnostic regular-expression matching (grep) of Ensembl identifiers ENSG00000120217, ENSG00000188389, and ENSG00000163599, respectively. The same barcode-based matching and risk-group assignments were used for the checkpoint analysis, comprising 197 observations (99 Low-Risk and 98 High-Risk) across 190 unique patients. Expression distributions were compared using two-sided Wilcoxon rank-sum tests, and nominal (unadjusted) p-values were visualized in Supplementary Figure 7.

### 2.5. Machine Learning Ensemble and Explainable Artificial Intelligence (AI)

Predictive modeling and feature prioritization were performed in Python (v3.10.12) using scikit-learn (v1.3.0–1.6.1) [16], imbalanced-learn (v0.12.0–0.14.1), xgboost, lightgbm (v4.0.0–4.6.0) [17], and shap (v0.44.0–0.51.0) [18]. Analyses used 190 deduplicated TCGA-BRCA TNBC patients with complete VST-normalized 47-gene expression and Vital_Status encoded as Alive = 0 (n = 163) or Dead = 1 (n = 27); random_state=42 was used for stochastic procedures. To generate the internal test data, the 47-gene matrix of 190 patients was first standardized using StandardScaler and subsequently balanced by the Synthetic Minority Over-sampling Technique (SMOTE) [19]. This process generated 136 synthetic Dead observations, resulting in 326 total observations (163 Alive, 163 Dead), which were then split 80%/20% to form the 66-observation test set. Because multi-omic discovery (Section 2.3), standardization, and SMOTE preceded the split, the test partition was not isolated from pre-split feature discovery, scaling, or synthetic interpolation; internal metrics therefore reflect performance within the SMOTE-augmented feature space.

Seven classifiers: Support Vector Machine (probability=True), Random Forest (n_estimators=300) [20], Gradient Boosting (n_estimators=200) [21], XGBoost (eval_metric=’logloss’), LightGBM [17], AdaBoost (n_estimators=100), and Logistic Regression (max_iter=2000)—were trained on 260 observations and evaluated on the 66-observation test set by accuracy and area under the receiver operating characteristic (ROC) curve (ROC-AUC) (Table 2). Random Forest Gini importance, corroborated by LightGBM, prioritized NKAIN3, VSX1, MEIS1, SLC1A4, FOXE1, SLC7A3, GCM2, CACNA1B, LRRC14, ECEL1, GAD2, PHACTR1, EN2, PCSK2, and TLX2 from the 47 genes (Figure 1). Random Forest and LightGBM were re-trained on this 15-gene panel using the same preprocessing, resampling, and partition. Pairwise Pearson correlations among the 15 genes and Vital_Status were calculated across the 190 unresampled patients (Supplementary Figure 5).

**Figure 1.**
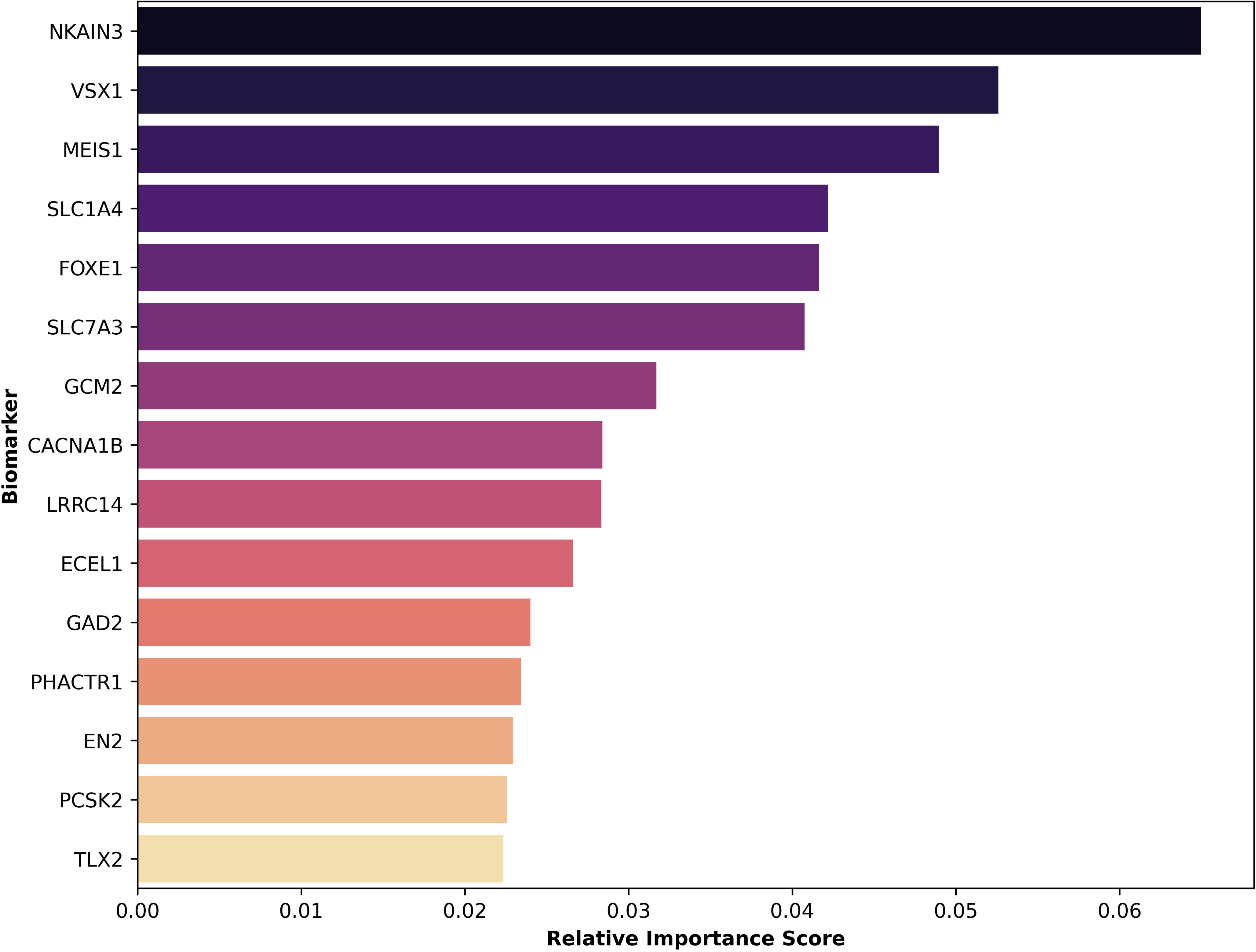
Machine-learning feature-importance ranking of 15 genes prioritized from the 47-gene discovery signature. Feature-importance analysis using Random Forest (corroborated with LightGBM) fitted on the internal training partition (n = 260 training observations from the n = 326 SMOTE-resampled TCGA-BRCA TNBC dataset) was used to prioritize the top 15 genes from the 47-gene discovery signature. The plot shows the relative Gini feature-importance scores of the selected genes. NKAIN3 showed the highest relative importance, followed by VSX1 and MEIS1, whereas TLX2 ranked 15th. The resulting 15 genes constituted the prognostic panel used for subsequent predictive and prognostic analyses.

**Table 2.** Internal predictive performance of seven machine-learning classifiers on the SMOTE-resampled TCGA-BRCA test partition.

| **Model** | **Correct Test Predictions (n / 66)** | **Accuracy (%)** | **ROC-AUC** |
|---|---|---|---|
| LightGBM | 63/66 | 95.45454545 | 0.989861751 |
| Support Vector Machine | 62/66 | 93.93939394 | 0.993548387 |
| Random Forest | 62/66 | 93.93939394 | 0.987096774 |
| Gradient Boosting | 60/66 | 90.90909091 | 0.982488479 |
| AdaBoost | 60/66 | 90.90909091 | 0.946543779 |
| XGBoost | 57/66 | 86.36363636 | 0.973271889 |
| Logistic Regression | 54/66 | 81.81818182 | 0.822119816 |

For explainable AI, LightGBM was fitted to all 326 SMOTE-resampled observations and tree-based SHapley Additive exPlanations (SHAP) values were computed using shap.TreeExplainer [18]. For Class 1 (Dead), positive values indicated contributions toward Dead predictions and negative values toward Alive predictions; the 15 features with the highest mean absolute SHAP values were displayed in Supplementary Figure 4.

To generate the final 190 patient-level ML Risk Scores, the following sequential pipeline was applied to the 15-gene panel: First, the 190-patient panel was rank-normalized to a standard Gaussian distribution using QuantileTransformer. Second, the data were balanced using SMOTE to generate 326 observations. Third, an equal-weight (1:1:1) soft-voting ensemble (LightGBM, Random Forest, and L2-penalized Logistic Regression) was fitted to the resampled data using five-fold Platt sigmoid calibration [22]. Finally, the averaged calibrated Class-1 probabilities (0–1) were extracted for the 190 original patients to serve as the continuous 15-gene ML Risk Score.

### 2.6. Time-Dependent Survival Analysis and Nomogram Construction

Overall survival (OS) was constructed in R by matching the patient-level 15-gene ML Risk Scores (n = 190) with the TCGA clinical (clinical.tsv) and follow-up (follow_up.tsv) datasets using the TCGA patient identifier (cases.submitter_id). Unavailable-data placeholders were converted to missing values before analysis. To ensure exactly one record per patient, repeated longitudinal records from the follow-up dataset were consolidated by calculating the maximum recorded follow-up duration for each patient. Vital status and days to death were obtained from the clinical dataset, while the maximum recorded follow-up duration was determined for each patient from the follow-up dataset. Time zero was the initial pathologic diagnosis, and OS was measured in days. Event status was coded as 1 for deceased patients using demographic.days_to_death and 0 for living patients right-censored at their maximum follow_ups.days_to_follow_up. Of the 190 patients with ML risk scores, three living patients had non-positive maximum follow-up durations and were excluded, leaving 187 patients (160 Alive and 27 Dead) for Kaplan–Meier and time-dependent ROC analyses.

Kaplan–Meier analysis [23] using the survival [24] and survminer packages stratified the 187-patient eligible cohort at the patient-level median ML Risk Score (approximately 0.006256) into Low-Risk (n = 94; Risk_Score ≤ 0.006256) and High-Risk (n = 93; Risk_Score > 0.006256) groups. Survival distributions were compared using the log-rank test. For multivariable Cox proportional hazards analysis [25], the ML Risk Score was standardized as a continuous Z-score and modeled with age per decade and AJCC pathologic stage (advanced III–IV versus early I–II reference) using survival [24]. The primary AJCC pathologic stage was selected from the diagnosis record designated as the primary disease for each patient. Complete-case filtering excluded four patients with missing or unclassifiable primary AJCC pathologic stage from the 187-patient OS cohort, leaving 183 patients (158 Alive and 25 Dead), comprising 154 patients with early-Stage disease and 29 with advanced Stage disease. Formal Schoenfeld residual testing of the proportional hazards assumption was not performed.

Time-dependent ROC analysis [26] using timeROC (v0.4) evaluated the ML Risk Score at 1, 3, and 5 years (365, 1,095, and 1,825 days) using the 187-patient OS cohort. A clinical-genomic nomogram [27] was constructed using rms (v8.1-1) [28] and a Cox model incorporating the original unstandardized ML Risk Score (0–1), age, and primary AJCC pathologic stage as a four-level factor in the 183-patient complete-case cohort (Stage I: n = 25; Stage II: n = 129; Stage III: n = 26; Stage IV: n = 3). Predictor points were summed and mapped to estimated 3- and 5-year OS probabilities.

### 2.7. Independent External Validation and Cross-Platform Evaluation

The 15-gene genomic risk signal was evaluated in the independent GSE58812 microarray cohort (n = 107 TNBC patients; 78 alive [72.9%], 29 deceased [27.1%]; Supplementary Table 4) on a platform distinct from TCGA-BRCA RNA-seq [2,11]. Oligonucleotide microarray expression profiles (Affymetrix Human Genome U133 Plus 2.0 Array; GSE58812_series_matrix.txt; GPL570) were extracted in R with GEOquery and mapped from probe sets to official gene symbols using hgu133plus2.db, retaining the first mapping for multi-mapped probes; unannotated probes were removed and duplicate symbols were resolved by retaining the first-mapped probe. All 47 discovery genes, including all 15 panel genes, were present without imputation. TCGA data comprised log2-scale VST-normalized RNA-seq counts, whereas GSE58812 values were retained as native linear microarray intensities without additional log2 transformation; consequently, applying the TCGA-fitted StandardScaler directly to unlogged microarray intensities introduced a cross-platform scale mismatch. A StandardScaler fitted to the 15-gene TCGA matrix was applied to GSE58812, and mortality was encoded as Alive = 0 and Dead = 1.

External evaluation comprised three stages. The first stage constituted the untouched independent external validation and was completed before any GSE58812-guided model optimization. In this stage, six fixed classifiers—Random Forest (n_estimators=500, class_weight=’balanced’) [20], XGBoost (eval_metric=’logloss’), LightGBM (is_unbalance=True) [17], Support Vector Machine (radial basis function kernel, probability=True, class_weight=’balanced’), Gradient Boosting (n_estimators=200) [21], and Logistic Regression (class_weight=’balanced’, max_iter=1000)—were trained on the SMOTE-resampled TCGA 15-gene data (n = 326) and evaluated directly on GSE58812 by ROC-AUC; random_state=42 was used for all six classifiers.

Second, after examination of the initial benchmark, XGBoost underwent post-validation Bayesian optimization with Optuna (30 trials; complete trial logs provided in supplementary materials) to maximize GSE58812 ROC-AUC over n_estimators 100–500, max_depth 3–10, learning_rate 0.01–0.3 (log-uniform), subsample 0.5–1.0, and colsample_bytree 0.5–1.0, with random_state=42 and eval_metric=’logloss’ fixed. Each candidate was trained on the n = 326 TCGA data and evaluated on GSE58812. The best estimator was calibrated using five-fold Platt sigmoid calibration [22] on the TCGA data and evaluated on GSE58812; significance of the calibrated gene-only ROC-AUC was assessed by a five-fold, 100-permutation ROC-AUC test.

Third, because pathological nodal stage was unavailable, age at diagnosis from Supplementary Table 4 (n = 107; mean = 56.97 years, sample standard deviation (SD) = 12.80, population SD = 12.74, range = 28.48–84.63) was Z-standardized within GSE58812. The standardized age Z-score was combined with calibrated genomic risk using a prespecified 0.7/0.3 genomic-risk-to-age weighting clipped to [0,1] to generate an exploratory genomic risk-plus-age score (Figure 6). In the no-nodal-covariate check, multiplying the combined score by 0.8 preserved patient rankings and ROC-AUC.

## Results

### 3.1. Baseline Patient Characteristics

The uncollapsed source TCGA-BRCA clinical table (clinical.tsv) comprised 5,546 treatment-level records from 1,098 unique patients (Table 1). Mean age at index was 56.71 ± 12.76 years (n = 5,545); among records with known vital status, 4,697 (84.7%) were Alive and 848 (15.3%) were Dead, and mean overall survival was 1,340.83 ± 1,208.26 days (n = 5,540). Among 4,978 records with available AJCC stage, Stage II (57.2%) and Stage III (25.4%) predominated.

Filtering for ER-negative, PR-negative, and HER2-negative primary solid tumors with PAM50 Basal classification and matched multi-omic data yielded 197 primary-tumor RNA-seq aliquots (168 Alive, 29 Dead) from 195 unique tumor specimens and 190 unique TNBC patients (Supplementary Table 1); these observations constituted the cohort for immune-infiltration and checkpoint analyses (Section 3.6). After deduplication of five patients with replicate aliquots, 190 patients with complete 47-gene measurements and known vital status were retained for machine-learning and SHAP analyses (163 Alive [85.8%], 27 Dead [14.2%]; mean age 55.36 ± 12.23 years; AJCC Stage I: n = 23, Stage II: n = 119, Stage III: n = 20, Stage IV: n = 3, Stage X/unreported: n = 25).

Linking the 190 patient-level ML Risk Scores with the TCGA clinical and follow-up datasets established the overall survival analysis cohort. Following exclusion of three living patients with non-positive maximum follow-up durations, 187 patients (160 Alive and 27 Dead) were retained for Kaplan–Meier (Section 3.5) and time-dependent ROC analyses (Section 3.7). Further complete-case filtering for missing or unclassifiable primary AJCC pathologic stage excluded four patients, resulting in a cohort of 183 patients (158 Alive and 25 Dead) for multivariable Cox regression (Section 3.5) and clinical-genomic nomogram construction (Section 3.7).

This table summarizes all 5546 uncollapsed treatment-level clinical rows in the source GDC TCGA-BRCA clinical.tsv table, representing 1098 unique patients; individual patients may contribute multiple rows. Reported characteristics include the number of clinical-record rows and unique patients, mean age at index with standard deviation (56.71 ± 12.76 years; n = 5545 non-missing records), AJCC pathologic tumor-stage distribution among 4978 records with non-missing stage (568 unreported), vital status among 5545 non-missing records (Alive: 4697 [84.7%]; Dead: 848 [15.3%]; 1 unreported), and overall survival time (1340.83 ± 1208.26 days; n = 5540 non-missing records). Percentages are calculated at the clinical-record level rather than the unique-patient level.

### 3.2. Multi-Omic Profiling and Initial Discovery

Principal component analysis (PCA) of the VST-transformed TCGA-BRCA transcriptomic matrix visualized sample-level variation across annotated sample types, with Solid Tissue Normal samples occupying a distinct cluster separated from Primary Tumor and Metastatic samples along Principal Component 1 and Principal Component 2 (Supplementary Figure 1). Single-factor DESeq2 differential expression analysis contrasting Primary Tumor (numerator) against Solid Tissue Normal reference samples (denominator) across 60,660 Ensembl gene identifiers (56,820 with evaluable test statistics; log2FC range: −6.43 to +6.31) identified 5,097 transcriptomic features with nominal p < 0.05 (1,651 upregulated [log2FC > 0] and 3,446 downregulated [log2FC < 0]; 551 with Benjamini–Hochberg padj < 0.05); at the volcano-plot visualization thresholds (|log2FC| > 1 and nominal p < 0.05), 326 genes were upregulated and 1,720 genes were downregulated (Supplementary Figure 2, Supplementary Table 2).

Parallel single-factor limma differential methylation analysis across the 10,000 filtered Illumina 450K CpG probes contrasting Primary Tumor against Solid Tissue Normal reference samples (logFC range: −0.21 to +0.23; AveExpr range: 0.0873–0.9163) identified 571 CpG probes with nominal P.Value < 0.05 (450 hypermethylated [logFC > 0] and 121 hypomethylated [logFC < 0]); no CpG probe reached Benjamini–Hochberg adj.P.Val < 0.05 (minimum adj.P.Val = 0.8709; Supplementary Table 3).

Separately from the tumor-versus-normal comparisons described above, the 47-gene discovery signature was derived from differential analyses based on patient vital status. RNA-seq expression differences between Dead and Alive patients were assessed using DESeq2, while DNA methylation differences between the same vital-status groups were assessed using limma. Genes with nominal p < 0.05 in the RNA-seq analysis and genes annotated to CpG probes with nominal P.Value < 0.05 in the methylation analysis were mapped to gene symbols and intersected. This procedure identified 47 overlapping candidate genes, each meeting the nominal significance criterion in both data types (Section 2.3). The resulting 47-gene signature was carried forward for functional enrichment and subsequent machine-learning feature prioritization.

### 3.3. Functional Enrichment and SHAP Interpretability

Hypergeometric functional enrichment analysis of the 47-gene discovery signature in Enrichr identified enriched biological pathways across the Reactome 2024, KEGG 2026, and WikiPathways 2024 libraries (Supplementary Figure 3). The top 10 ranked pathways from each database encompassed metabolic, signaling, developmental, solute-transport, neuronal, and differentiation-related processes. TreeExplainer SHAP analysis of the LightGBM classifier fitted to the 326 SMOTE-resampled 47-gene observations characterized the magnitude and directionality of individual gene contributions toward mortality prediction (Class 1 = Dead; Supplementary Figure 4). The SHAP summary plot displays the 15 features with the highest mean absolute SHAP values from the 47-gene discovery set, where positive SHAP values drive predictions toward Dead and negative SHAP values drive predictions toward Alive.

### 3.4. Selection of the 15-Gene Prognostic Panel

Across the 190 unique TNBC patients, StandardScaler standardization and SMOTE resampling balanced the binary outcome classes (163 Alive, 163 Dead; n = 326), and the resulting dataset was partitioned into 260 training observations (80%; 128 Alive, 132 Dead) and 66 internal test observations (20%; 35 Alive, 31 Dead). Table 2 summarizes the internal classification accuracy and ROC-AUC of the seven evaluated machine-learning classifiers fitted on the n = 260 training partition and evaluated on the held-out n = 66 test partition. LightGBM achieved the highest classification accuracy (95.45% [63/66]; ROC-AUC = 0.9899), whereas Support Vector Machine achieved the highest ROC-AUC (0.9935; accuracy = 93.94% [62/66]), followed by Random Forest (accuracy = 93.94% [62/66]; ROC-AUC = 0.9871), Gradient Boosting (accuracy = 90.91% [60/66]; ROC-AUC = 0.9825), AdaBoost (accuracy = 90.91% [60/66]; ROC-AUC = 0.9465), XGBoost (accuracy = 86.36% [57/66]; ROC-AUC = 0.9733), and Logistic Regression (accuracy = 81.82% [54/66]; ROC-AUC = 0.8221).

Feature-importance ranking from the Random Forest model fitted on the n = 260 training partition (corroborated with LightGBM) prioritized the top 15 prognostic biomarkers from the 47-gene signature (NKAIN3, VSX1, MEIS1, SLC1A4, FOXE1, SLC7A3, GCM2, CACNA1B, LRRC14, ECEL1, GAD2, PHACTR1, EN2, PCSK2, and TLX2; Figure 1), showing a hierarchical importance distribution led by NKAIN3, VSX1, and MEIS1, with TLX2 ranking 15th. Re-training Random Forest and LightGBM on the prioritized 15-gene matrix (TCGA_TNBC_Top15_ML_Dataset.csv) under the identical pre-split StandardScaler, SMOTE, and 80%/20% partition (n = 260 train; n = 66 test) confirmed high internal discrimination (Random Forest accuracy = 93.94% [62/66], ROC-AUC = 0.9880; LightGBM ROC-AUC = 0.99). Pairwise Pearson correlation coefficients among the 15 prioritized genes and numerically encoded Vital_Status across the 190 unresampled unique patients were visualized in Supplementary Figure 5.

The 5-fold sigmoid-calibrated soft-voting ensemble (LightGBM, Random Forest, and Logistic Regression) subsequently generated the continuous 15-gene ML Risk Score (TCGA_TNBC_Top15_Risk_Scores.csv; overall range: 0.000111–0.998196; mean = 0.1577 ± 0.3445; median = 0.006083; interquartile range [IQR] = 0.002355–0.021089) for all 190 unique TNBC patients (Alive [n = 163]: mean = 0.0269 ± 0.1104, median = 0.005227; Dead [n = 27]: mean = 0.9476 ± 0.1775, median = 0.988189).

Predictive performance is presented as correct test predictions, classification 466 accuracy (%), and area under the receiver operating characteristic curve (ROC-AUC) on the 20% internal test partition (n = 66 observations), derived from an 80%/20% random train/test split (n = 260 train, n = 66 test) of the SMOTE-resampled TCGA-BRCA TNBC cohort (n = 326 balanced observations generated from n = 190 unique TNBC patients: 163 Alive and 27 Dead). The evaluated models were LightGBM, Support Vector Machine, Random Forest, Gradient Boosting, AdaBoost, XGBoost, and Logistic Regression. LightGBM achieved the highest classification accuracy (95.45%), whereas Support Vector Machine achieved the highest ROC-AUC (0.9935). Because multi-omic discovery, standardization, and SMOTE preceded the train/test split, the test partition was not isolated from pre-split processing; these metrics therefore reflect internal performance within the SMOTE-augmented feature space.

### 3.5. Kaplan–Meier Survival and Multivariable Cox Regression

Among the 187 patients with eligible overall survival data, Kaplan–Meier analysis of the 15-gene ML Risk Score demonstrated significant separation between the Low-Risk (n = 94) and High-Risk (n = 93) groups, defined using the patient-level median risk score of approximately 0.006256. The High-Risk group showed poorer overall survival than the Low-Risk group (log-rank p < 0.0001; Figure 2).

**Figure 2.**
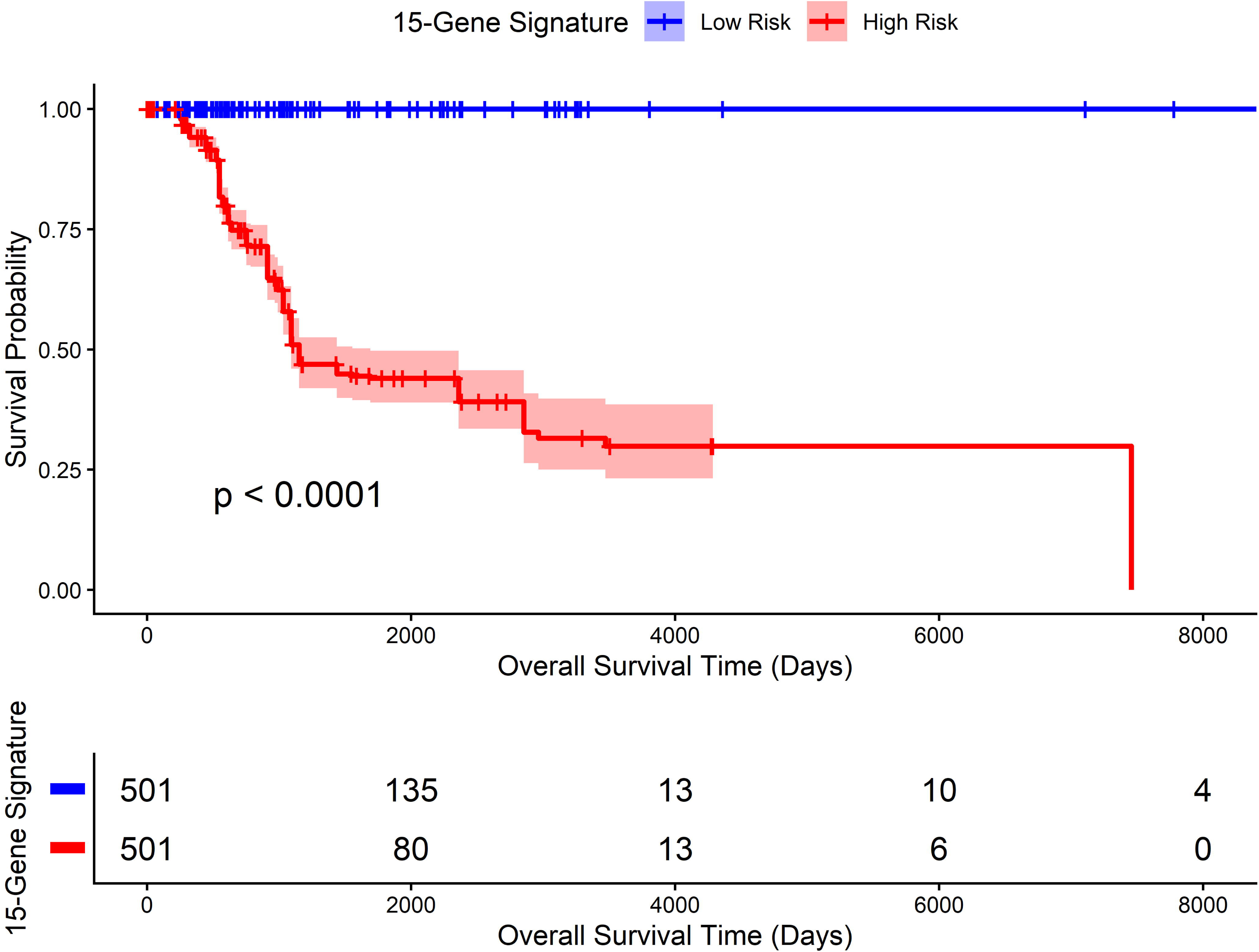
Kaplan–Meier curves for High-Risk and Low-Risk groups defined by the 15-gene ML Risk Score. The 187-patient eligible overall survival cohort was stratified by the patient-level median 15-gene ML Risk Score (approximately 0.006256) into Low-Risk (n = 94) and High-Risk (n = 93) groups. Overall survival was evaluated using Kaplan–Meier analysis, and survival distributions were compared using the log-rank test. The High-Risk group showed poorer overall survival than the Low-Risk group (log-rank p < 0.0001).

To evaluate the prognostic association of the ML Risk Score after adjustment for clinical covariates, multivariable Cox proportional hazards regression was performed in the 183-patient complete-case cohort. The model incorporated the standardized continuous ML Risk Score, age per decade, and AJCC pathologic stage categorized as advanced (III–IV) versus early (I–II). The cohort comprised 158 Alive and 25 Dead patients, including 154 patients with early-stage disease and 29 with advanced-stage disease. A one-standard-deviation increase in the ML Risk Score was associated with a hazard ratio of 10.67 (95% confidence interval (CI): 6.64–17.15; p < 0.001). Advanced AJCC stage was also associated with increased hazard (hazard ratio (HR) = 3.76; 95% CI: 2.57–5.51; p < 0.001), whereas age per decade was not significantly associated with hazard (HR = 0.90; 95% CI: 0.75–1.07; p = 0.226; Figure 3, Table 3).

**Figure 3.**
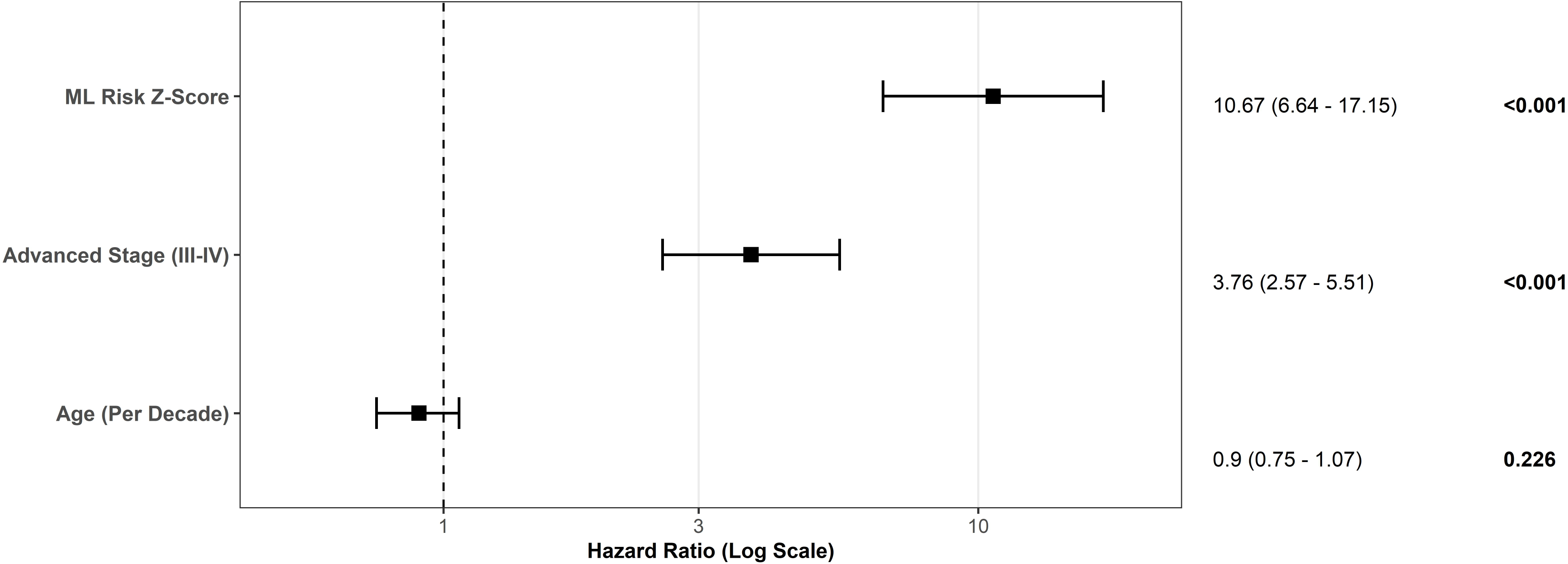
Multivariable Cox proportional hazards analysis of the 15-gene ML Risk Score (Z-score) and clinical covariates. The multivariable Cox proportional hazards model was fitted in the 183-patient complete-case cohort and included the standardized continuous 15-gene ML Risk Score, patient age per decade, and AJCC pathologic stage categorized as advanced (III–IV) versus early (I–II reference). Hazard ratios (HRs) and 95% confidence intervals (CIs) are shown. A one-standard-deviation increase in the ML Risk Score was associated with a hazard ratio of 10.67 for overall survival (95% CI, 6.64–17.15; p < 0.001). Advanced AJCC pathologic stage was also associated with increased hazard (HR, 3.76; 95% CI, 2.57–5.51; p < 0.001), whereas age, modeled per decade, was not significantly associated with hazard (HR, 0.90; 95% CI, 0.75–1.07; p = 0.226).

**Table 3.** Multivariable Cox regression of overall survival with 15-gene ML Risk Score and clinical covariates.

| **Variable** | **HR** | **95% CI** | **P value** |
|---|---|---|---|
| **Age (per decade)** | 0.90 | 0.75 - 1.07 | 0.226 |
| **Advanced AJCC pathologic stage (III–IV vs. I–II)** | 3.76 | 2.57 - 5.51 | <0.001 |
| **15-gene ML Risk Score (Z-score)** | 10.67 | 6.64 - 17.15 | <0.001 |

The multivariable Cox proportional hazards model was fitted in the 183-patient complete-case cohort (158 Alive and 25 Dead), comprising 154 patients with early-stage disease and 29 with advanced-stage disease. The model evaluated the association of the 15-gene ML Risk Score with overall survival after adjustment for patient age and primary AJCC pathologic stage, categorized as advanced (III–IV) versus early (I–II reference). The ML Risk Score was standardized as a continuous Z-score; therefore, its hazard ratio represents the change in hazard associated with a one-standard-deviation increase in the original ML Risk Score. Age was modeled per decade. Hazard ratios (HRs), 95% confidence intervals (CIs), and P values are presented for each covariate. The 15-gene ML Risk Score (HR, 10.67; 95% CI, 6.64–17.15; P < 0.001) and advanced AJCC pathologic stage (HR, 3.76; 95% CI, 2.57–5.51; P < 0.001) were significantly associated with overall survival, whereas age was not significantly associated with hazard (HR, 0.90; 95% CI, 0.75–1.07; P = 0.226).

### 3.6. Immune Microenvironment and Checkpoint Analysis

The ssGSEA analysis characterized estimated immune-infiltration scores across the six-population canonical immune panel in the matched TCGA-BRCA TNBC cohort (n = 197 primary-tumor observations across 190 unique patients; n = 99 Low-Risk and n = 98 High-Risk defined by the median 15-gene ML Risk Score of 0.005983164; Supplementary Figure 6). Two-sided Wilcoxon rank-sum tests across all six evaluated immune profiles (visualized in Supplementary Figure 6) showed no statistically significant differences between the Low-Risk and High-Risk cohorts: CD8+ T cells (Low-Risk median = 0.2498 [interquartile range, IQR = 0.1918] vs. High-Risk median = 0.2414 [IQR = 0.1730], p = 0.93), B cells (median = 0.1890 [IQR = 0.2280] vs. 0.1650 [IQR = 0.2180], p = 0.68), M1 macrophages (median = 0.3784 [IQR = 0.1646] vs. 0.3662 [IQR = 0.1400], p = 0.91), M2 macrophages (median = 0.2837 [IQR = 0.1313] vs. 0.2760 [IQR = 0.1530], p = 0.91), regulatory T cells (median = 0.2210 [IQR = 0.1267] vs. 0.2324 [IQR = 0.1324], p = 0.38), and natural killer (NK) cells (median = 0.2213 [IQR = 0.1165] vs. 0.2051 [IQR = 0.1027], p = 0.53). These findings indicate that the 15-gene ML Risk Score was not significantly associated with any of the six ssGSEA immune-infiltration profiles evaluated in Supplementary Figure 6.

Separately, we compared VST-normalized targeted immune checkpoint gene expression between the same median-defined Low-Risk (n = 99) and High-Risk (n = 98) cohorts. No statistically significant differences were observed for CD274/PD-L1 (p = 0.72), PDCD1/PD-1 (p = 0.74), or CTLA4 (p = 0.50) (Supplementary Figure 7).

### 3.7. Time-Dependent ROC Analysis and Clinical-Genomic Nomogram Construction

Time-dependent ROC analysis evaluated the prognostic discrimination of the 15-gene ML Risk Score in the 187-patient overall survival cohort at 1, 3, and 5 years (Figure 4). The corresponding time-specific AUCs were 0.910, 0.911, and 0.937, respectively. A clinical-genomic nomogram was constructed in the 183-patient complete-case cohort by integrating the original unstandardized 15-gene ML Risk Score (0–1), patient age, and primary AJCC pathologic stage as a four-level categorical predictor (Stage I: n = 25; Stage II: n = 129; Stage III: n = 26; Stage IV: n = 3). Predictor points were summed to generate a total score, which was mapped to estimated 3-year and 5-year overall survival probabilities (Figure 5).

**Figure 4.**
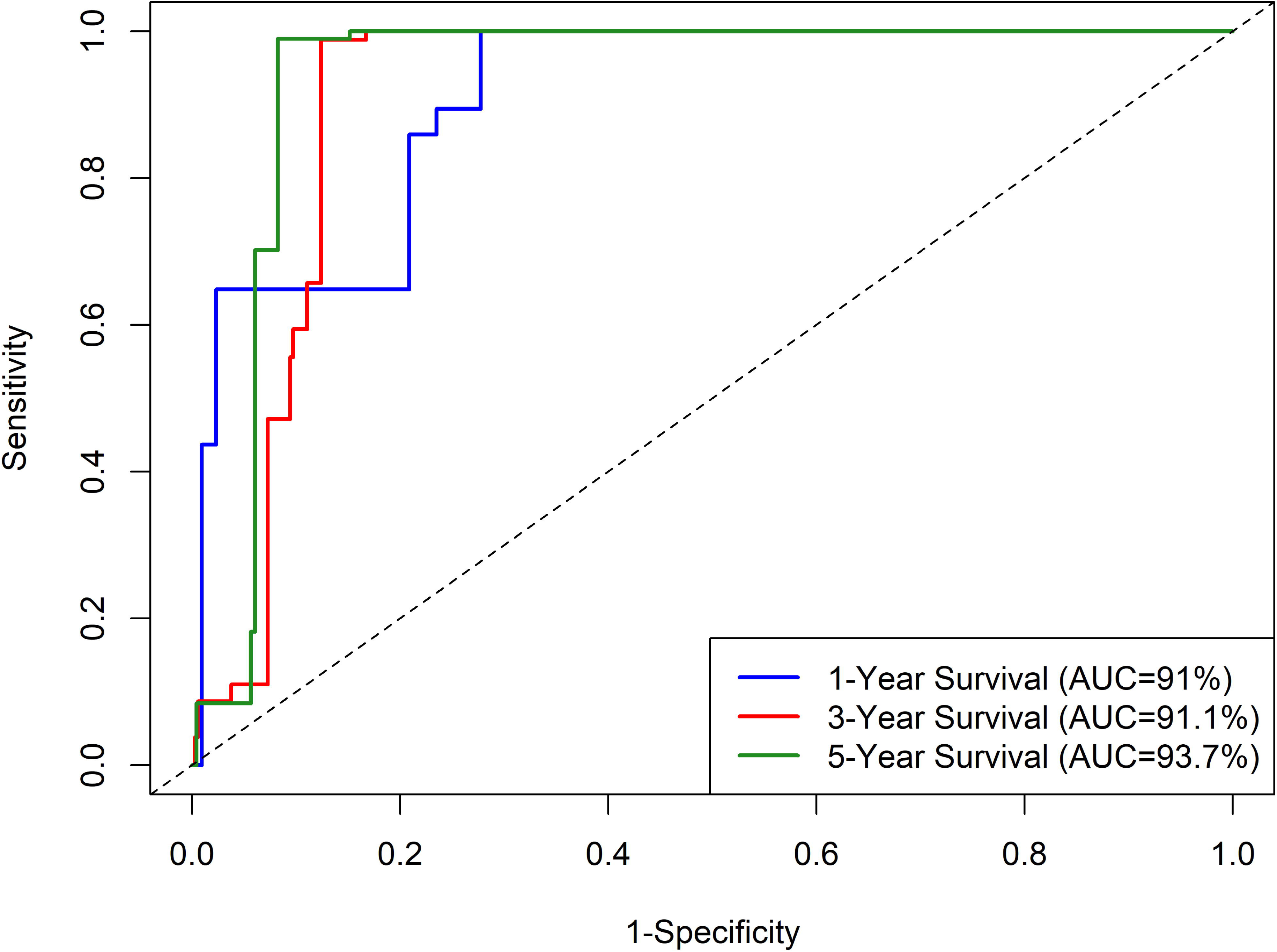
Time-dependent ROC analysis of 15-gene ML Risk Score for 1-, 3-, and 5-year overall survival. Time-dependent receiver operating characteristic (ROC) analysis was performed in the 187-patient eligible overall survival cohort to assess the prognostic discrimination of the 15-gene ML Risk Score at 1, 3, and 5 years. The corresponding time-specific areas under the ROC curves (AUCs) were 0.910 at 1 year, 0.911 at 3 years, and 0.937 at 5 years. Sensitivity is plotted against 1-specificity for each evaluated time point.

**Figure 5.**
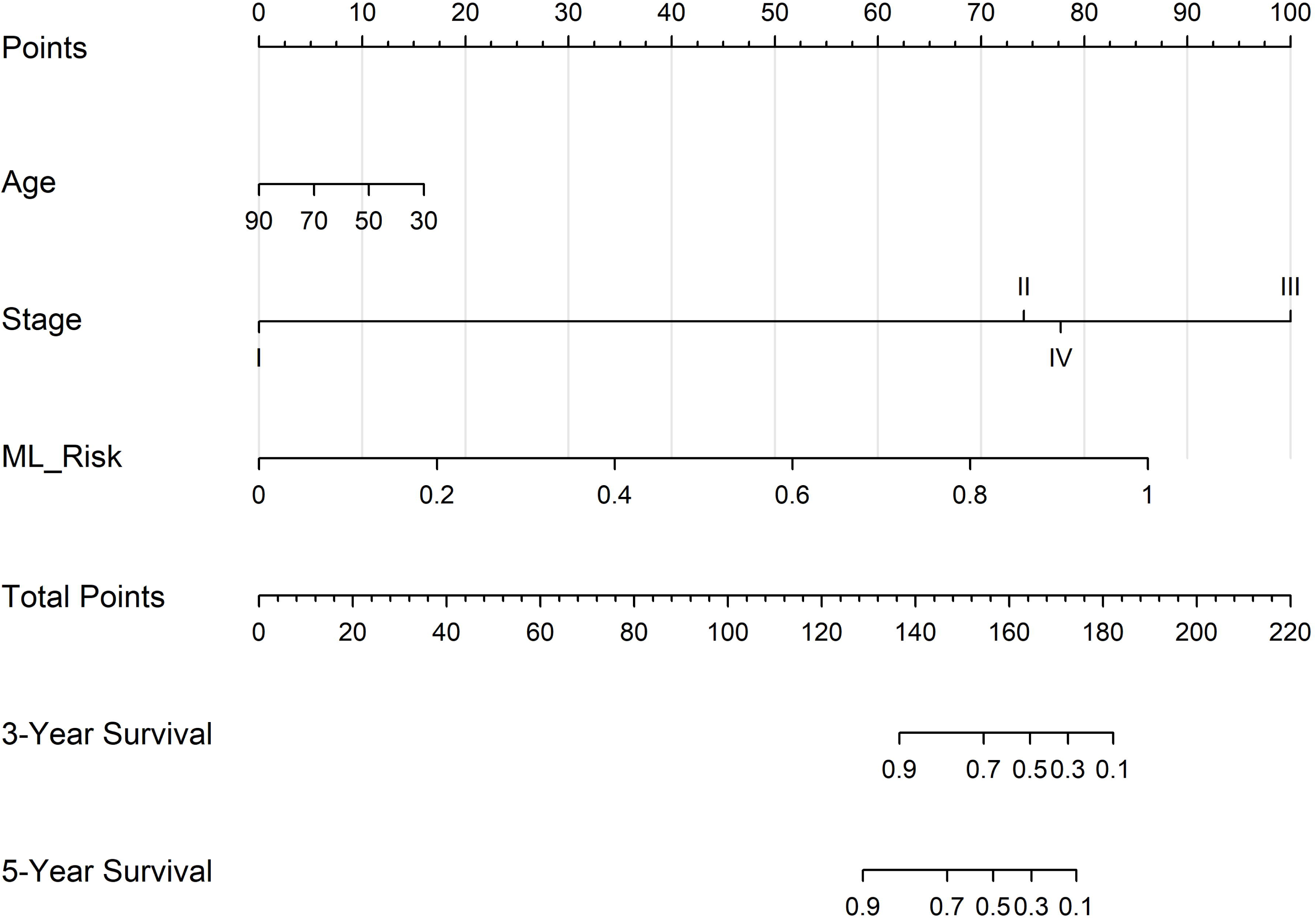
Clinical-genomic nomogram for 3- and 5-year survival: age, AJCC stage, and 15-gene ML Risk Score. The clinical-genomic nomogram was constructed in the 183-patient complete-case cohort and incorporated the original unstandardized 15-gene ML Risk Score (0–1), patient age, and primary AJCC pathologic stage modeled as a four-level categorical predictor (Stage I: n = 25; Stage II: n = 129; Stage III: n = 26; Stage IV: n = 3). Predictor points were summed to obtain a total point score, which was mapped to estimated 3-year and 5-year overall survival probabilities.

**Figure 6.**
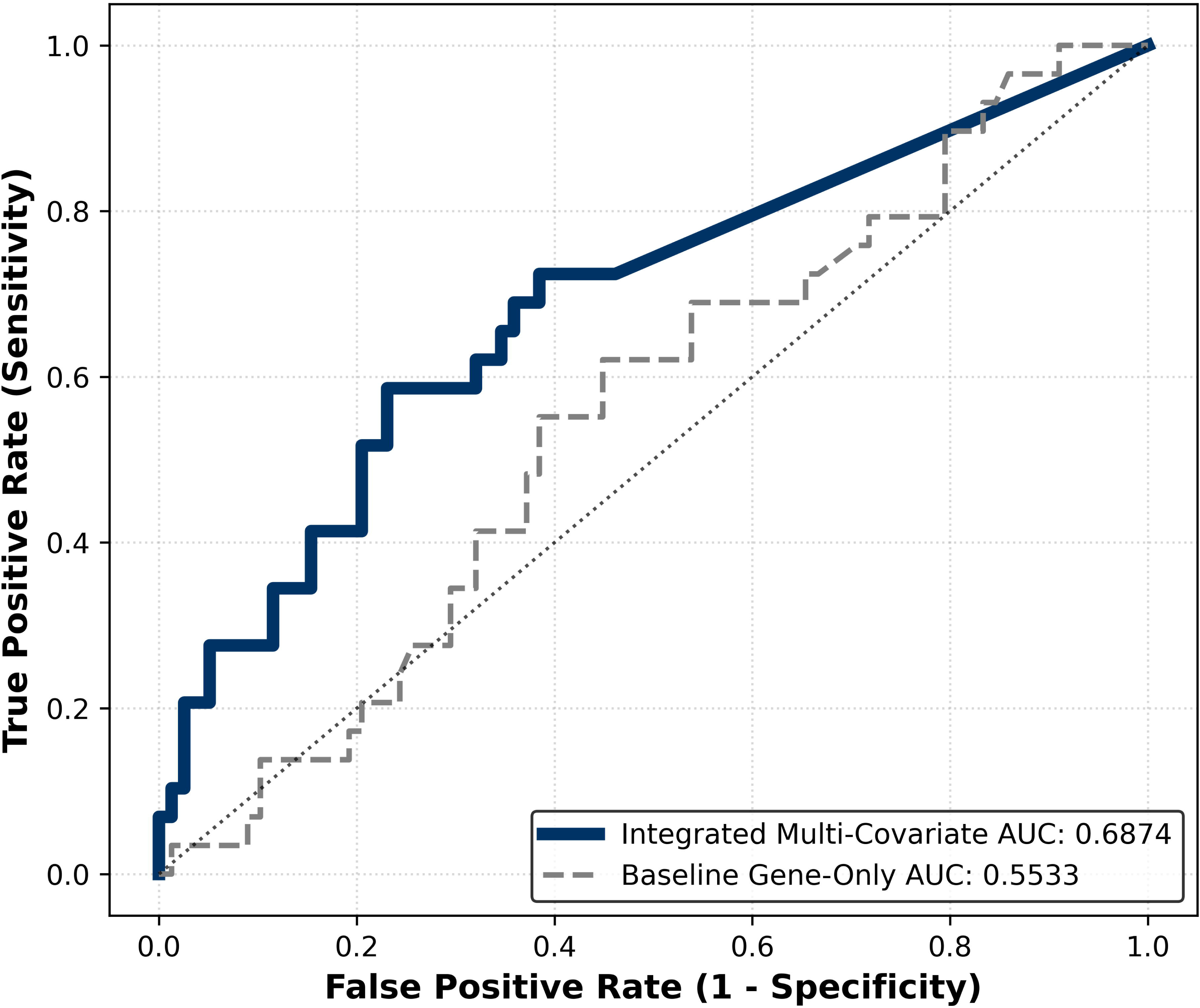
External validation and subsequent analysis of the 15-gene genomic risk signature in GSE58812. In the independent GSE58812 cohort (n = 107), the initial untouched six-model external-validation benchmark yielded ROC-AUC values of 0.4191–0.5500. The curves shown represent subsequent analyses: post-validation optimized and calibrated XGBoost yielded a gene-only ROC-AUC of 0.5533, while an exploratory genomic risk-plus-age score yielded an ROC-AUC of 0.6874. These subsequent results should not be interpreted as untouched external-validation performance. The ROC curves show sensitivity versus 1-specificity.

### 3.8. Independent External Validation and Predictive Performance

All 47 discovery genes, including all 15 prioritized panel genes, were successfully mapped without imputation in the independent GSE58812 Affymetrix Human Genome U133 Plus 2.0 microarray cohort (n = 107 TNBC patients; 78 alive [72.9%], 29 deceased [27.1%]). The external cohort therefore contained the complete genomic feature set required for cross-platform evaluation. However, the TCGA-BRCA discovery cohort comprised log2-scale VST-normalized RNA-seq counts, whereas GSE58812 retained native linear microarray intensities and was transformed using the TCGA-fitted StandardScaler parameters; consequently, feature distributions were not represented in identical measurement units across the two platforms.

The initial six-model benchmark constituted the untouched independent external validation and was completed before any GSE58812-guided model optimization. In this benchmark, six fixed machine-learning classifiers trained on the SMOTE-resampled TCGA 15-gene data (n = 326) were evaluated directly on GSE58812 and showed limited discrimination, with ROC-AUC values ranging from 0.4191 to 0.5500. XGBoost yielded the highest ROC-AUC (0.5500 [0.549956]), followed by Random Forest (0.5442), Logistic Regression (0.5371), Support Vector Machine (0.5000), LightGBM (0.4478), and Gradient Boosting (0.4191). Thus, the high internal discrimination observed in the TCGA-derived analyses was not reproduced in the untouched independent cross-platform external-validation benchmark.

Following completion and examination of the independent external-validation benchmark, post-validation Bayesian optimization of XGBoost using Optuna (30 trials), followed by five-fold sigmoid probability calibration, yielded a gene-only ROC-AUC of 0.5533. This discrimination did not reach statistical significance in the 100-permutation test (p = 0.9208). Because pathological nodal stage was unavailable in GSE58812, patient age at diagnosis (56.97 ± 12.80 years) was evaluated as an available clinical covariate. Combining Z-score standardized age with calibrated genomic risk using the prespecified 0.7/0.3 genomic-risk-to-age weighting yielded an exploratory integrated genomic risk-plus-age ROC-AUC of 0.6874 (Figure 6). The optimized gene-only and age-integrated results represent post-validation and exploratory analyses, respectively, rather than the untouched independent external-validation performance of the fixed models.

## Discussion

Targeted therapeutic options for TNBC remain limited, reinforcing the need for reproducible prognostic biomarkers and clinically interpretable risk models. In this study, we integrated transcriptomic and DNA-methylation information to derive a 47-gene discovery signature and used Random Forest and LightGBM feature-importance analyses to prioritize a focused 15-gene panel across 190 unique TCGA-BRCA TNBC patients. Internally, LightGBM achieved the highest classification accuracy (95.45%; ROC-AUC = 0.9899), whereas Support Vector Machine achieved the highest ROC-AUC (0.9935; accuracy = 93.94%) on the SMOTE-resampled test partition. Functional enrichment provided biological context for the discovery signature, while SHAP analysis characterized the contribution and directionality of individual features relative to model output [18]. These findings are consistent with previous TNBC studies demonstrating the value of integrating complementary molecular layers [4,5] and applying computational feature selection to reduce high-dimensional molecular data to smaller prognostic signatures [6,7]. Chappell et al. identified correlated regulatory features across multiple molecular layers in TNBC [4], Hashem and Mabrouk integrated transcriptomic and DNA-methylation data to identify molecular alterations associated with TNBC metabolism and survival [5], and machine-learning studies by Kim et al. and Hwangbo et al. similarly developed focused prognostic models from high-dimensional molecular data [6,7]. The 15-gene panel therefore provides a focused candidate set for further assessment using targeted expression assays such as reverse transcription quantitative polymerase chain reaction (RT-qPCR), although its analytical performance and clinical utility remain to be established through experimental and independent clinical validation.

The 15-gene ML Risk Score also showed prognostic association within the TCGA-derived survival analyses. Kaplan–Meier analysis of the 187-patient eligible overall survival cohort showed significant separation between the Low-Risk and High-Risk groups. In the 183-patient complete-case cohort, multivariable Cox regression demonstrated that the standardized ML Risk Score remained associated with overall survival after adjustment for patient age and AJCC pathologic stage. A one-standard-deviation increase in the ML Risk Score was associated with a hazard ratio of 10.67 (95% CI: 6.64–17.15; p < 0.001), while advanced stage was also associated with increased hazard (HR = 3.76; 95% CI: 2.57–5.51; p < 0.001). Time-dependent AUCs were 0.910, 0.911, and 0.937 at 1, 3, and 5 years, respectively. The clinical-genomic nomogram integrated the original unstandardized ML Risk Score with patient age and primary AJCC pathologic stage to estimate 3-year and 5-year overall survival probabilities. Such nomograms can combine multiple predictors into individualized outcome estimates, although the statistical uncertainty and validation status of those estimates must be considered [27].

Immune-infiltration and immune-checkpoint analyses provided exploratory biological context but did not identify statistically significant differences between the ML-defined risk groups. Across 197 matched primary-tumor observations from 190 unique patients, none of the six evaluated ssGSEA immune-infiltration profiles differed significantly between Low-Risk and High-Risk cohorts (p = 0.38–0.93), and no significant differences were observed for CD274/PD-L1, PDCD1/PD-1, or CTLA4 expression. These findings indicate that the prognostic stratification captured by the 15-gene ML Risk Score was not significantly associated with the specific immune variables evaluated here. This does not exclude broader immune heterogeneity in TNBC: Jézéquel et al. identified molecular subgroups with distinct immune-response and macrophage characteristics [2], while Han et al. described a cytotoxic-lymphocyte-related signature associated with survival and immune-checkpoint features [8]. Rather, the present findings indicate that the six selected immune profiles and three checkpoint genes did not distinguish the risk groups defined by this genomic score.

The independent cross-platform evaluation of the TCGA-derived 15-gene genomic signal in GSE58812 represents an important methodological strength relative to studies relying exclusively on internal validation. The untouched benchmark of six fixed classifiers, completed before any GSE58812-guided optimization, yielded ROC-AUC values of 0.4191–0.5500, demonstrating substantial attenuation relative to the internal results. Subsequent post-validation optimization of XGBoost followed by five-fold sigmoid calibration yielded a gene-only AUC of 0.5533, with no evidence of statistical significance in the permutation test (p = 0.9208), whereas the exploratory genomic risk-plus-age score yielded an AUC of 0.6874. Because XGBoost optimization was conducted after examination of the initial GSE58812 results, these later estimates should be interpreted as post-validation, hypothesis-generating analyses rather than as untouched external validation of a prespecified model. This distinction is particularly important when comparing the present study with reports such as Hwangbo et al., in which selected TNBC predictive markers were evaluated across five independent datasets [7].

Several limitations should be considered. First, this retrospective computational study used publicly available datasets, and the internal patient-level machine-learning analysis included 190 unique TNBC patients. Multi-omic candidate discovery, feature scaling, and SMOTE oversampling were performed before the 80%/20% train/test split, and the calibrated voting ensemble was fitted on the resampled discovery cohort; therefore, the internal test partition was not isolated from pre-split discovery, scaling, or synthetic interpolation, and the internal predictive and survival estimates are subject to optimism from pre-split processing and in-sample risk-score calibration. Second, survival analyses were based on patient-level matching of the 15-gene ML Risk Scores with the TCGA clinical and follow-up datasets. The eligible overall survival cohort comprised 187 patients for Kaplan–Meier and time-dependent ROC analyses, while complete-case filtering for missing or unclassifiable primary AJCC pathologic stage yielded 183 patients for multivariable Cox regression and nomogram construction. Third, the immune-infiltration and immune-checkpoint analyses retained replicate aliquots from five patients, resulting in 197 aliquot-level observations from 190 unique patients. The Wilcoxon rank-sum tests treated these observations as independent and did not explicitly account for within-patient correlation among replicate aliquots. In addition, nominal, unadjusted p-values were reported without multiple-testing correction.

Fourth, GSE58812 was generated on an Affymetrix microarray platform using linear probe intensities, whereas TCGA used log2-VST RNA-seq data; applying TCGA-fitted StandardScaler parameters directly to unlogged microarray intensities introduced cross-platform scale mismatch and technical variation that contributed to the attenuation of external discrimination. Fifth, the subsequent XGBoost optimization was post-validation, and the genomic risk-plus-age score was exploratory rather than a jointly fitted clinical-genomic model because age was combined post hoc with genomic risk probabilities using the prespecified 0.7/0.3 weighting. Finally, formal proportional-hazards diagnostics were not performed, the nomogram was developed without internal bootstrap calibration or independent external validation, and the 15-gene panel has not undergone experimental or prospective clinical validation. These considerations reflect broader challenges in translating machine-learning models in oncology, including data heterogeneity, bias, interpretability, and the distinction between retrospective proof-of-concept analyses and independently validated models [10].

Overall, this study provides a computational framework linking multi-omic discovery, machine-learning feature prioritization, prognostic modeling, selected immune-context analysis, and independent cross-platform external validation in TNBC. The 15-gene panel showed significant prognostic association within the TCGA-derived analyses, whereas the independent GSE58812 evaluation demonstrated limited gene-only transferability across platforms. The post-validation age-integrated analysis suggests that clinical information may provide additional signal, but this finding remains exploratory. Further evaluation in larger independent cohorts with standardized cross-platform processing [29], prospectively defined clinical-genomic modeling, and experimental validation of the selected biomarkers will be required before clinical implementation [9,30].

## Conclusions

By integrating transcriptomic and epigenomic discovery with machine-learning feature prioritization, prognostic modeling, and selected immune-context analyses, we reduced a 47-gene discovery signature to a focused 15-gene prognostic panel. The 15-gene ML Risk Score demonstrated significant survival separation between the Low-Risk and High-Risk groups in the 187-patient eligible overall survival cohort. In the 183-patient complete-case cohort, the ML Risk Score remained associated with overall survival after adjustment for patient age and AJCC pathologic stage, with a hazard ratio of 10.67 per one-standard-deviation increase. Time-dependent ROC analysis showed AUCs of 0.910, 0.911, and 0.937 at 1, 3, and 5 years, respectively. The clinical-genomic nomogram integrated the original ML Risk Score with patient age and primary AJCC pathologic stage to estimate 3-year and 5-year overall survival probabilities, although its performance requires independent validation. Immune microenvironment and checkpoint analyses showed no statistically significant differences between High-Risk and Low-Risk cohorts for the evaluated ssGSEA immune-infiltration profiles or targeted immune checkpoint expression variables. These findings indicate that the prognostic signal was not significantly associated with the specific immune variables assessed in this study.

Importantly, the TCGA-derived 15-gene genomic signal underwent untouched independent cross-platform external validation in GSE58812 before any GSE58812-guided model optimization; the initial six fixed-model benchmark yielded AUC values ranging from 0.4191 to 0.5500, indicating limited gene-only transferability across platforms. Post-validation XGBoost optimization yielded a gene-only AUC of 0.5533, while an exploratory genomic risk-plus-age score yielded an AUC of 0.6874; these later results represent post-validation and exploratory analyses, respectively, rather than untouched external-validation performance, and require independent assessment.

Overall, this study provides a computational approach for narrowing multi-omic TNBC discovery findings to a focused candidate prognostic panel and evaluating its transferability in an independent cohort measured on a distinct expression platform. This independent cross-platform evaluation is methodologically important because it provides a more stringent assessment of signature transferability than internal performance alone and highlights the need for standardized independent validation before clinical translation. The findings support further assessment of the panel’s reproducibility and prognostic value in larger independent cohorts, with standardized cross-platform processing and experimental validation before any clinical implementation.

## Abbreviations

AI: Artificial Intelligence
AJCC: American Joint Committee on Cancer
AUC: area under the receiver operating characteristic curve
CI: confidence interval
CpG: cytosine-phosphate-guanine
DNA: deoxyribonucleic acid
ER: estrogen receptor
GDC: Genomic Data Commons
GEO: Gene Expression Omnibus
GSVA: Gene Set Variation Analysis
HER2: human epidermal growth factor receptor 2
HR: hazard ratio
IQR: interquartile range
KEGG: Kyoto Encyclopedia of Genes and Genomes
ML: machine learning
NCBI: National Center for Biotechnology Information
NK: natural killer
OS: overall survival
PAM50: Prediction Analysis of Microarray 50
PCA: principal component analysis
PD-1: programmed cell death protein 1
PD-L1: programmed death-ligand 1
PR: progesterone receptor
RNA-seq: RNA sequencing
ROC: receiver operating characteristic
ROC-AUC: area under the receiver operating characteristic curve
RT-qPCR: reverse transcription quantitative polymerase chain reaction
SD: standard deviation
SHAP: SHapley Additive exPlanations
SMOTE: Synthetic Minority Over-sampling Technique
ssGSEA: Single-Sample Gene Set Enrichment Analysis
TCGA: The Cancer Genome Atlas
TCGA-BRCA: The Cancer Genome Atlas breast cancer cohort
TNBC: triple-negative breast cancer
VST: variance-stabilizing transformation

## Declarations

### Ethics approval and consent to participate

This study involved the secondary analysis of publicly available, de-identified human genomic and clinical data from The Cancer Genome Atlas breast cancer cohort (TCGA-BRCA) and the independent GSE58812 external-validation cohort. The study did not involve the direct recruitment of human participants, collection of new biological samples, or access to personally identifiable information. Therefore, additional institutional ethics approval and informed consent were not required for this secondary analysis; accordingly, an ethics committee name and reference number were not applicable to the present study. The data were accessed and analyzed in accordance with the applicable data-access and data-use policies of the respective public repositories.

### Consent for publication

Not applicable.

### Availability of data and materials

The datasets used and/or analysed during the current study are publicly available from the Genomic Data Commons (GDC) Data Portal for the TCGA-BRCA project (https://portal.gdc.cancer.gov/projects/TCGA-BRCA) [1] and the National Center for Biotechnology Information (NCBI) Gene Expression Omnibus (GEO) for GSE58812, which served as the independent cross-platform external-validation cohort (https://www.ncbi.nlm.nih.gov/geo/query/acc.cgi?acc=GSE58812) [2,11]. Aggregated cohort composition and selected demographic, clinical, pathological-stage, and molecular-subtype characteristics of the TCGA-BRCA internal discovery cohort are provided in Supplementary Table 1. The complete DESeq2 differential-expression results and limma differential-methylation results are provided in Supplementary Tables 2 and 3, respectively. Aggregated clinical, demographic, diagnostic, vital-status, immunohistochemical, and platform characteristics of the independent GSE58812 external-validation cohort are provided in Supplementary Table 4.

### Authors’ contributions

N.M. conceptualised the study, developed the methodology, performed analyses, and drafted the manuscript. E.N. provided critical supervision, validated models, and reviewed the final manuscript.

## Acknowledgements

We thank the contributors to The Cancer Genome Atlas (TCGA) and the Gene Expression Omnibus (GEO) for making the publicly available datasets used in this study accessible to the research community.

## Authors’ information

N.M. and E.N. are affiliated with Biotecnika, Bengaluru, Karnataka, India.

---

**Supplementary Figure 1. Principal component analysis of VST-transformed transcriptomic data across annotated sample types.**

Exploratory principal component analysis (PCA) was performed using variance-stabilizing-transformed (DESeq2::vst) transcriptomic data from the TCGA-BRCA dataset to visualize global sample-level variation and distribution across all annotated TCGA sample types (Metastatic, Primary Tumor, and Solid Tissue Normal). The PCA plot displays the distribution of samples along Principal Component 1 and Principal Component 2, with Solid Tissue Normal samples largely occupying a distinct region separated from the Primary Tumor distribution. These three sample-type labels were used for exploratory PCA visualization only; downstream differential analyses contrasted Primary Tumor samples against Solid Tissue Normal reference samples.

**Supplementary Figure 2. Dual-threshold volcano plot of differential gene expression.**

Single-factor DESeq2 differential gene-expression results contrasting TCGA-BRCA Primary Tumor samples (numerator condition) against Solid Tissue Normal samples (reference denominator condition) across 56,820 evaluable Ensembl genes (from 60,660 total identifiers in Supplementary Table 2) are displayed as a dual-threshold volcano plot. The x-axis represents estimated log2 fold change (log2FC), and the y-axis represents statistical significance as −log10(nominal p-value). For visualization, transcriptomic features were classified as upregulated when log2FC > 1 and nominal p < 0.05 (n = 326, red), downregulated when log2FC < −1 and nominal p < 0.05 (n = 1,720, blue), and non-significant otherwise (grey). These results summarize the tumour-versus-normal differential-expression analysis described in Section 2.2 and are distinct from the vital-status-based selection of the 47-gene discovery signature described in Section 2.3.

**Supplementary Figure 3. Hypergeometric pathway enrichment of the 47-gene discovery signature across Reactome, KEGG, and WikiPathways.**

Hypergeometric functional enrichment analysis was performed for the 47-gene discovery signature using the Enrichr platform against the Reactome 2024, KEGG 2026, and WikiPathways 2024 libraries. The top 10 ranked pathways from each library are shown in panels (a), (b), and (c), respectively. The x-axis represents enrichment significance as −log10(p-value), with larger values indicating smaller enrichment p-values. The dashed vertical line represents the p = 0.05 reference threshold. Pathways are ranked according to enrichment p-value within each database.

**Supplementary Figure 4. SHAP summary plot of the 15 highest-ranked features from the 47-gene LightGBM model.**

SHapley Additive exPlanations (SHAP) analysis (shap.TreeExplainer) was performed using the LightGBM model fitted across the 326 SMOTE-resampled observations (derived from the 190 unique TCGA-BRCA TNBC patients) with the 47-gene discovery feature set to characterize the contribution and directionality of individual features relative to model output. The plot displays the 15 features with the highest mean absolute SHAP values (max_display = 15). The x-axis represents the SHAP value for Class 1 (Dead), with positive values indicating contributions toward the Dead prediction and negative values indicating contributions toward the Alive prediction. The color scale represents the corresponding standardized feature value, with red indicating higher feature values and blue indicating lower feature values.

**Supplementary Figure 5. Pearson correlation heatmap of 15 prioritized genes and numerically encoded Vital_Status in TNBC.**

Pairwise Pearson correlation analysis was performed across the 190 unresampled unique TCGA-BRCA TNBC patients (TCGA_TNBC_Top15_ML_Dataset.csv) among the 15 machine-learning-prioritized genes and the numerically encoded Vital_Status variable (Alive = 0 [n = 163] and Dead = 1 [n = 27]). The heatmap displays the Pearson correlation coefficients for each pair of variables, with blue indicating negative correlations, red indicating positive correlations, and diagonal cells representing self-correlations (r = 1.00).

**Supplementary Figure 6. Comparison of ssGSEA immune-infiltration scores between High-Risk and Low-Risk cohorts.**

Single-sample gene set enrichment analysis (ssGSEA) was performed using the GSVA package to estimate infiltration scores for six canonical immune cell populations in the matched TCGA-BRCA TNBC cohort (n = 197 primary-tumor observations across 190 unique patients). Observations were stratified into Low-Risk (n = 99) and High-Risk (n = 98) cohorts using the median 15-gene ML Risk Score (0.005983164) as the cutoff. Two-sided Wilcoxon rank-sum tests were used to compare ssGSEA immune-infiltration scores between the two risk groups across all six evaluated immune profiles: CD8+ T cells (p = 0.93), B cells (p = 0.68), M1 macrophages (p = 0.91), M2 macrophages (p = 0.91), regulatory T cells (p = 0.38), and NK cells (p = 0.53). None of the six evaluated profiles showed a statistically significant difference between the Low-Risk and High-Risk cohorts.

**Supplementary Figure 7. Comparison of VST-transformed expression of selected immune checkpoint genes between High-Risk and Low-Risk cohorts.**

VST-normalized expression of the immune checkpoint genes CD274 (PD-L1; ENSG00000120217), PDCD1 (PD-1; ENSG00000188389), and CTLA4 (ENSG00000163599) was compared between Low-Risk (n = 99) and High-Risk (n = 98) cohorts (n = 197 matched primary-tumor observations across 190 unique patients) defined using the median 15-gene ML Risk Score (0.005983164). Two-sided Wilcoxon rank-sum tests were used to compare expression distributions between the two risk groups. The boxplots show the distributions of normalized expression values, with individual observations displayed as points. The nominal (unadjusted) p-values were 0.72 for CD274 (PD-L1), 0.74 for PDCD1 (PD-1), and 0.50 for CTLA4. None of the three immune checkpoint genes showed a statistically significant difference in expression between the Low-Risk and High-Risk cohorts.

**Supplementary Table 1. Aggregated clinical and sample characteristics of the TCGA-BRCA TNBC discovery cohort.**

This table summarizes the aggregated clinical and sample characteristics of the TCGA-BRCA triple-negative breast cancer (TNBC) internal discovery cohort. The source dataset comprised 197 primary-tumor RNA-seq aliquots representing 195 unique tumor specimens and 190 unique patients whose tumors met the study eligibility criteria: estrogen receptor-negative, progesterone receptor-negative, human epidermal growth factor receptor 2-negative, and PAM50 Basal. Five patients contributed replicate primary-tumor aliquots: three patients contributed two aliquots each, and two patients contributed three aliquots each. All 197 aliquots were retained for immune-infiltration and immune-checkpoint analyses, whereas replicate aliquots were deduplicated at the patient level for machine-learning classification, SHAP interpretation, and risk-score generation (n = 190). Patient-level demographic and clinical characteristics are summarized using one record per unique patient. Percentages are calculated using the stated denominators, and missing or unreported categories are presented explicitly.

**Supplementary Table 2. Complete DESeq2 differential-expression results for the TCGA-BRCA transcriptomic dataset (60660 Ensembl gene identifiers).**

Complete gene-level differential-expression results were generated using negative binomial generalized linear models in DESeq2 (v1.38.3), contrasting Primary Tumor samples (numerator condition) against Solid Tissue Normal samples (reference denominator condition) in an unadjusted single-factor design (∼ sample_type) without additional clinical covariates. The table reports the Ensembl gene identifier, base mean expression (baseMean), estimated log2 fold change (log2FoldChange; range: −6.4283 to +6.3071), standard error (lfcSE), Wald test statistic (stat), nominal p-value (pvalue), and Benjamini–Hochberg adjusted p-value (padj). The supplementary table contains the complete unfiltered DESeq2 output: 60660 total rows, 56820 rows with non-missing test statistics, 30465 rows with non-missing padj, 5097 rows with nominal pvalue < 0.05 (1651 upregulated and 3446 downregulated), and 551 rows with padj < 0.05. These results summarize the tumour-versus-normal differential-expression analysis described in Section 2.2 and are distinct from the vital-status-based selection of the 47-gene discovery signature described in Section 2.3.

**Supplementary Table 3. Complete limma differential-methylation results for the 10000 analyzed Illumina 450K CpG probes.**

Complete differential-methylation results were generated using empirical Bayes moderated linear models (lmFit and eBayes) in limma (v3.54.2) across 10000 pre-processed Illumina HumanMethylation450 (450K) BeadChip CpG probes with complete beta-value measurements, contrasting Primary Tumor samples (numerator condition) against Solid Tissue Normal reference samples (denominator condition) in an unadjusted single-factor design (∼ sample_type) without additional covariates. The table reports the CpG probe identifier, limma coefficient (logFC; range: −0.2091 to +0.2307), average methylation beta-value (AveExpr; range: 0.0873 to 0.9163), moderated t-statistic (t), nominal P-value (P.Value), Benjamini–Hochberg adjusted P-value (adj.P.Val; minimum = 0.8709), and log-odds B-statistic (B). The supplementary table contains the complete 10000-probe limma output. These results summarize the tumour-versus-normal differential-methylation analysis described in Section 2.2 and are distinct from the vital-status-based selection of the 47-gene discovery signature described in Section 2.3. No probe reached adj.P.Val < 0.05.

**Supplementary Table 4. Clinical and demographic characteristics of the independent GSE58812 validation cohort.**

Aggregate clinical, demographic, diagnostic, and technical characteristics of the independent GSE58812 external validation cohort comprising 107 triple-negative breast cancer patients, each represented by one primary-tumor expression profile. Age at diagnosis is summarized as mean ± standard deviation and range. Vital status and metastatic-status source codes are reported as cohort-level counts and percentages. All 107 samples were annotated as triple-negative breast cancer with negative estrogen receptor, progesterone receptor, and HER2 immunohistochemistry. Expression profiling was performed using the Affymetrix Human Genome U133 Plus 2.0 Array (GPL570), with MAS 5.0 normalization. Individual sample identifiers and participant-level clinical and survival records are not included.

