## Supplementary figures and images for "Transcriptomic-Methylation Integration and Machine-Learning Prioritization of Candidate Prognostic Biomarkers in Triple-Negative Breast Cancer: A Retrospective Computational Study"

### Supplementary Figure 1

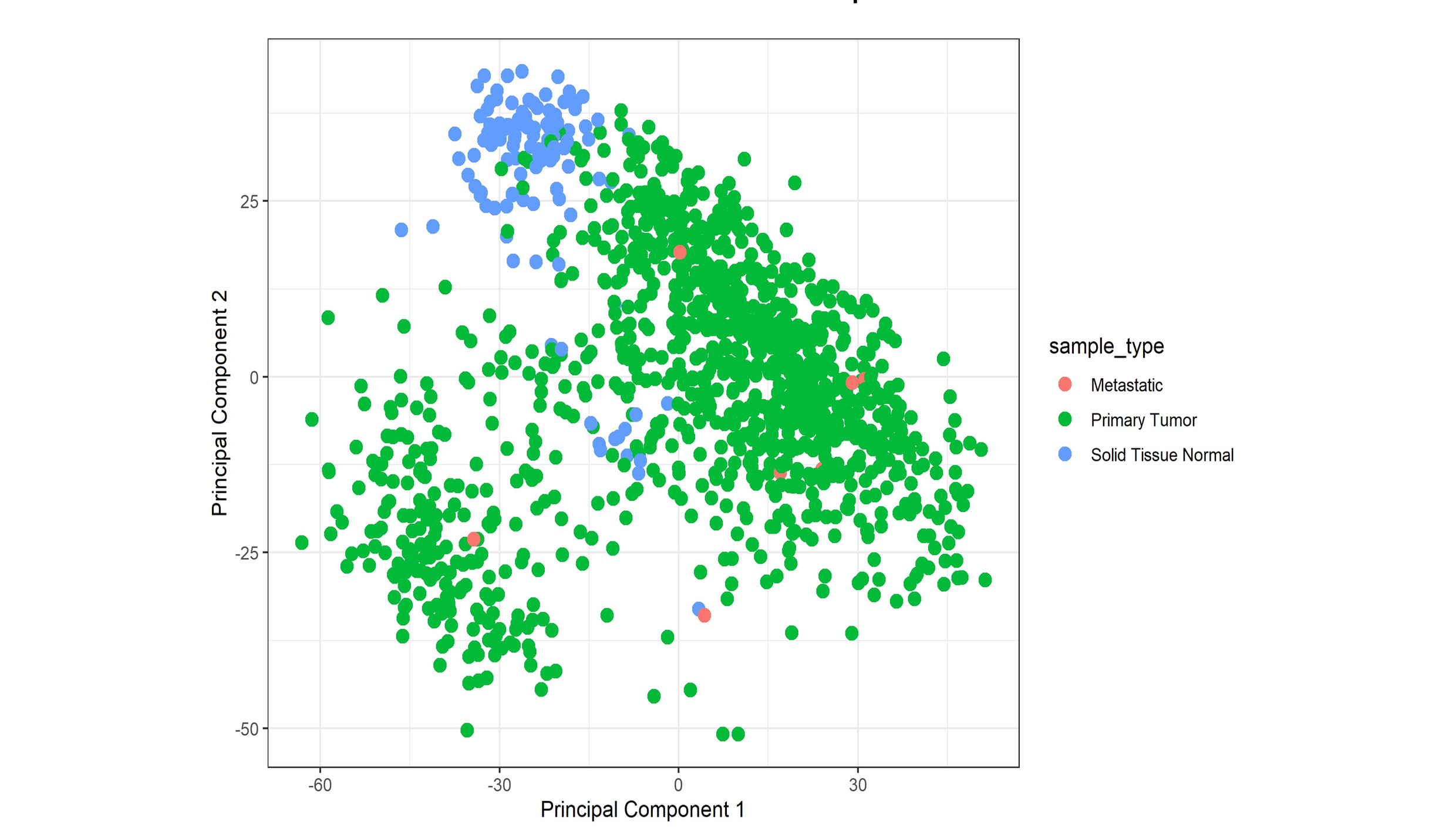

### Supplementary Figure 2

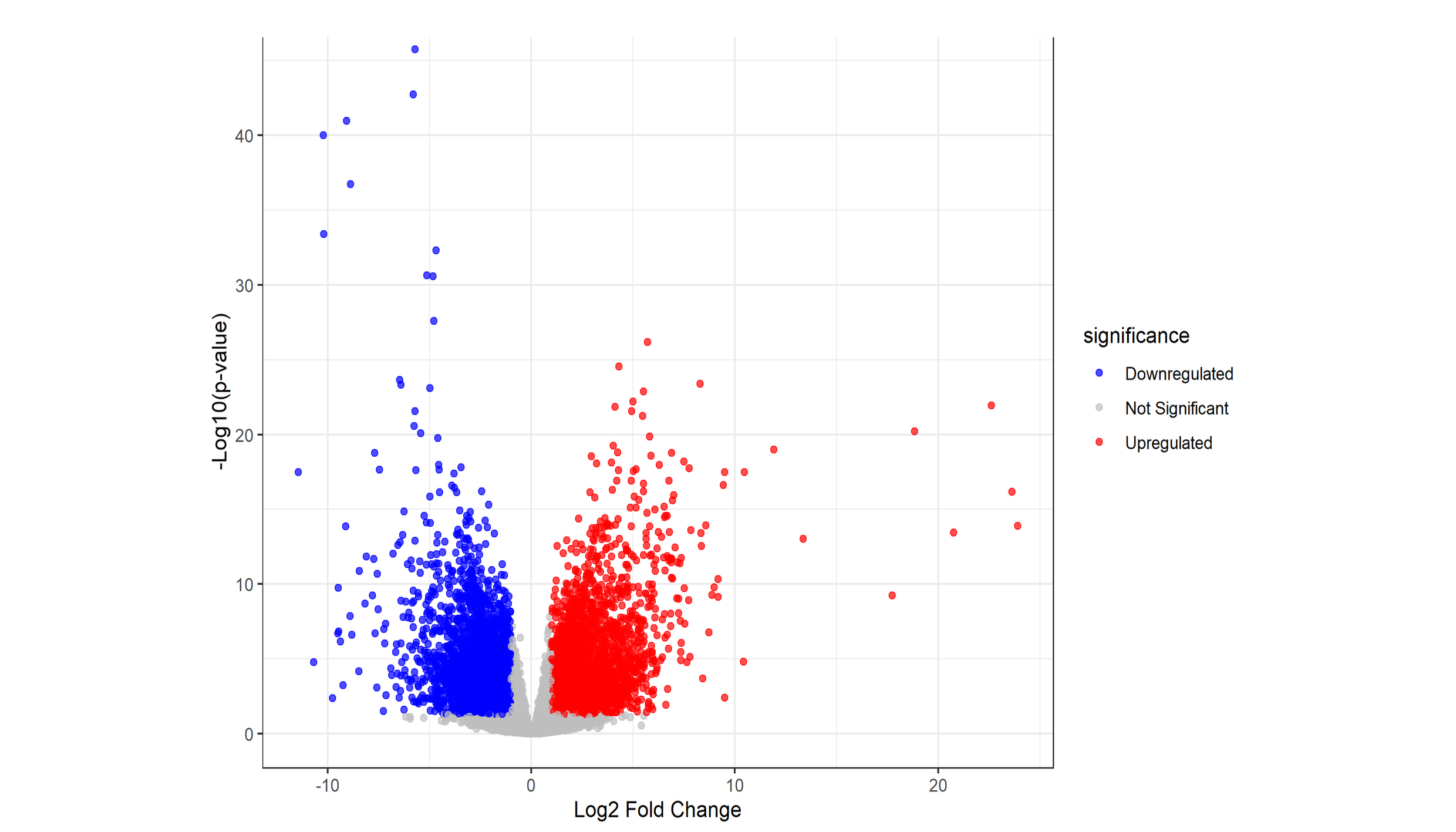

### Supplementary Figure 3

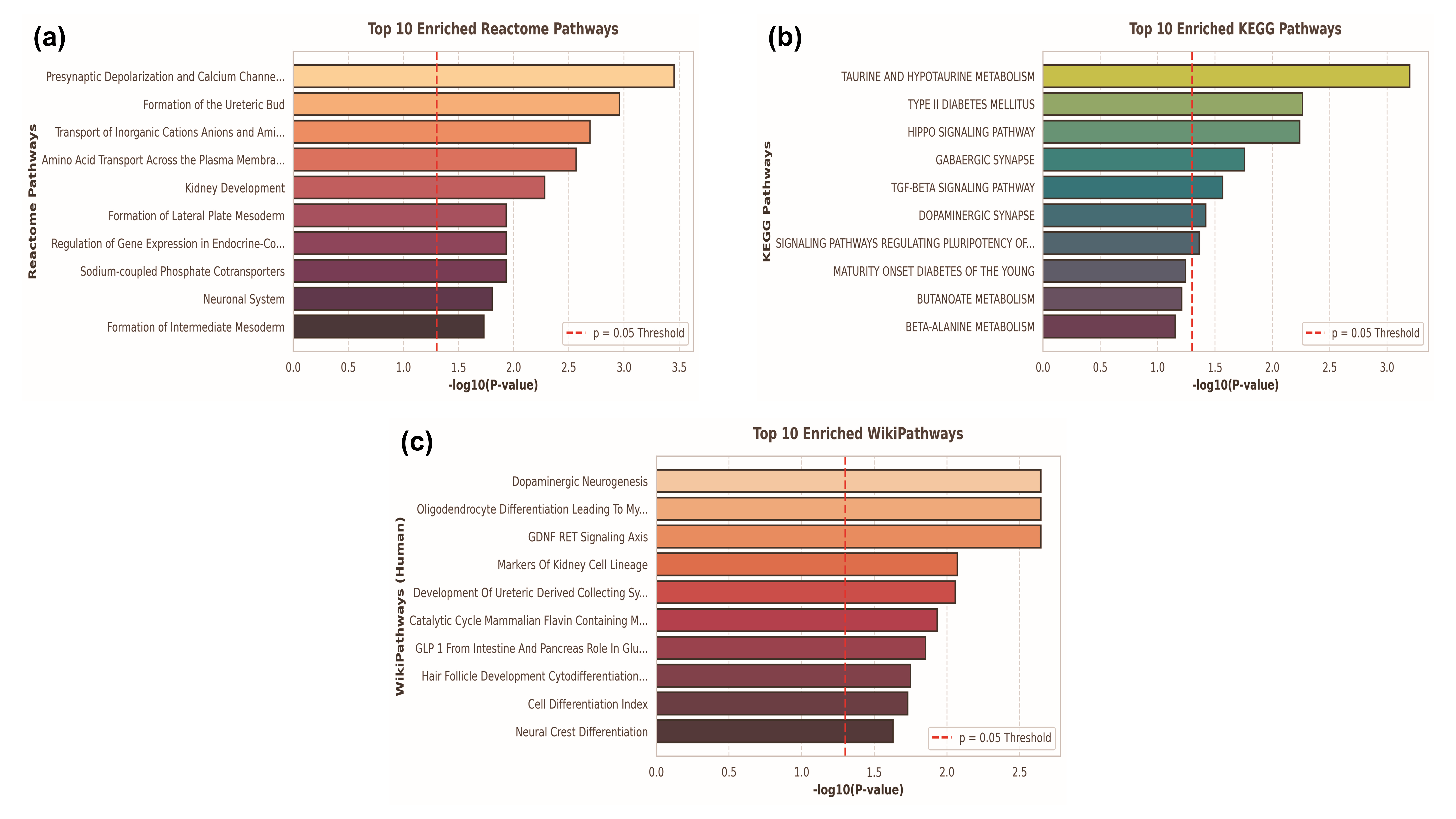

### Supplementary Figure 4

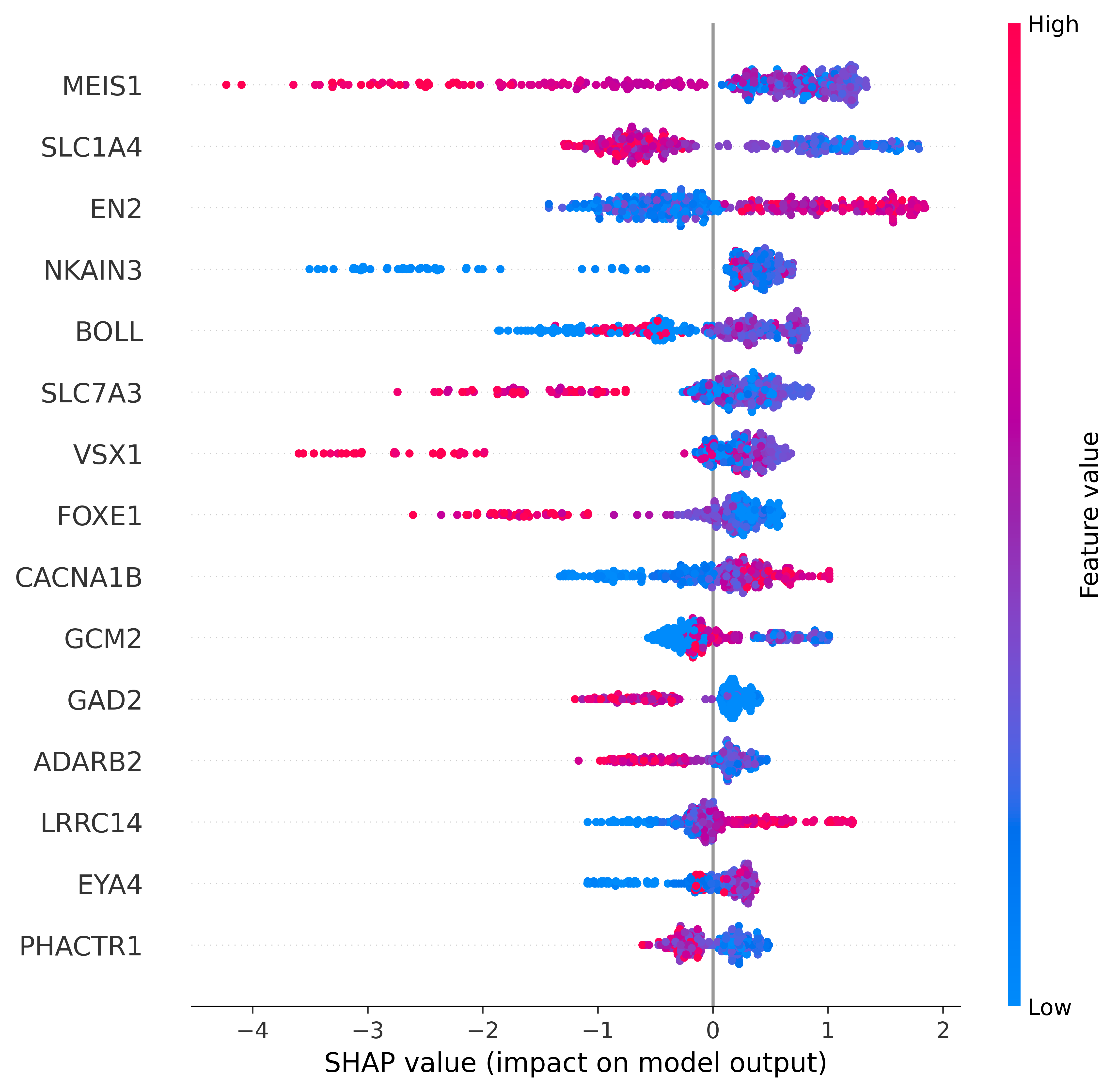

### Supplementary Figure 5

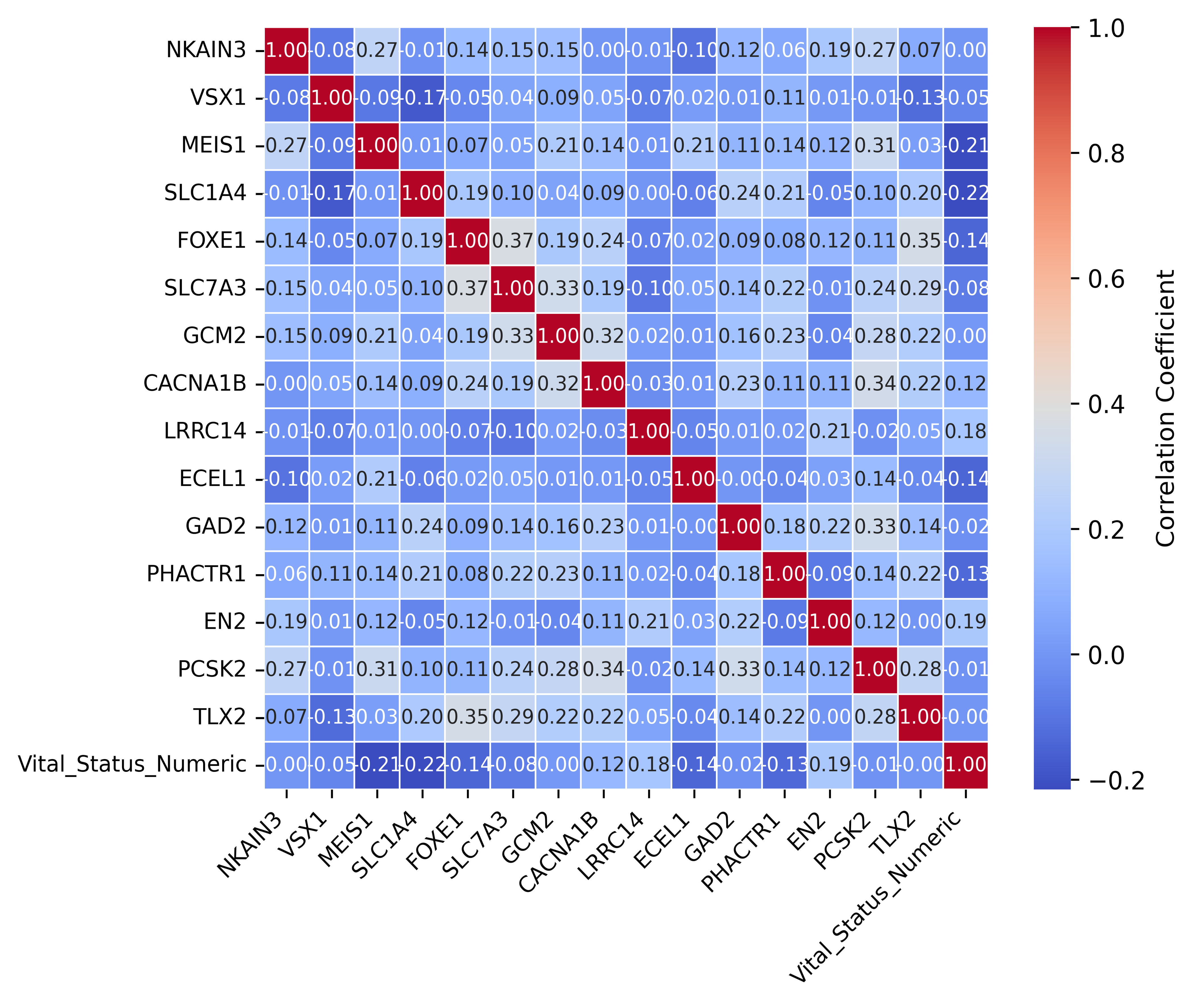

### Supplementary Figure 6

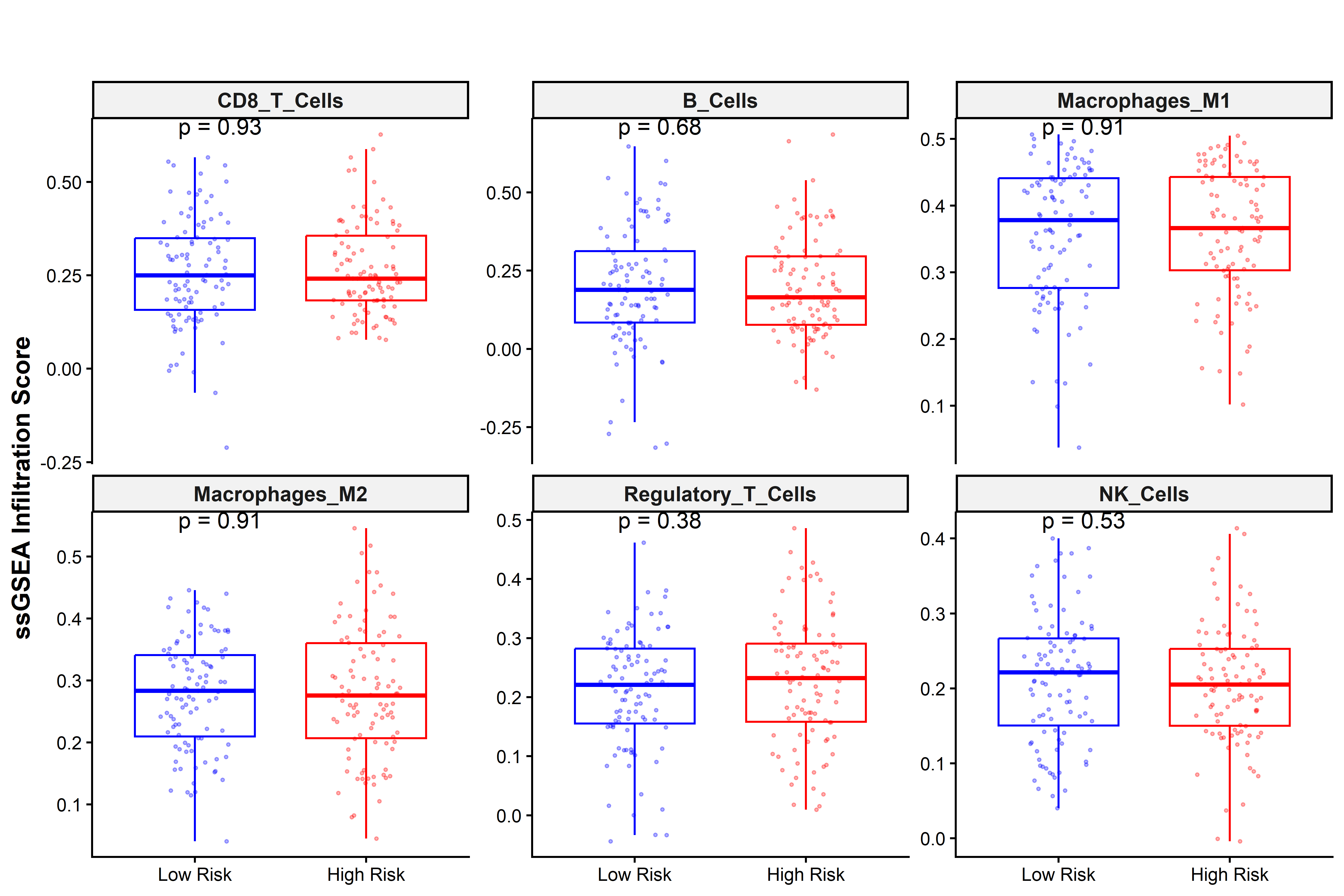

### Supplementary Figure 7

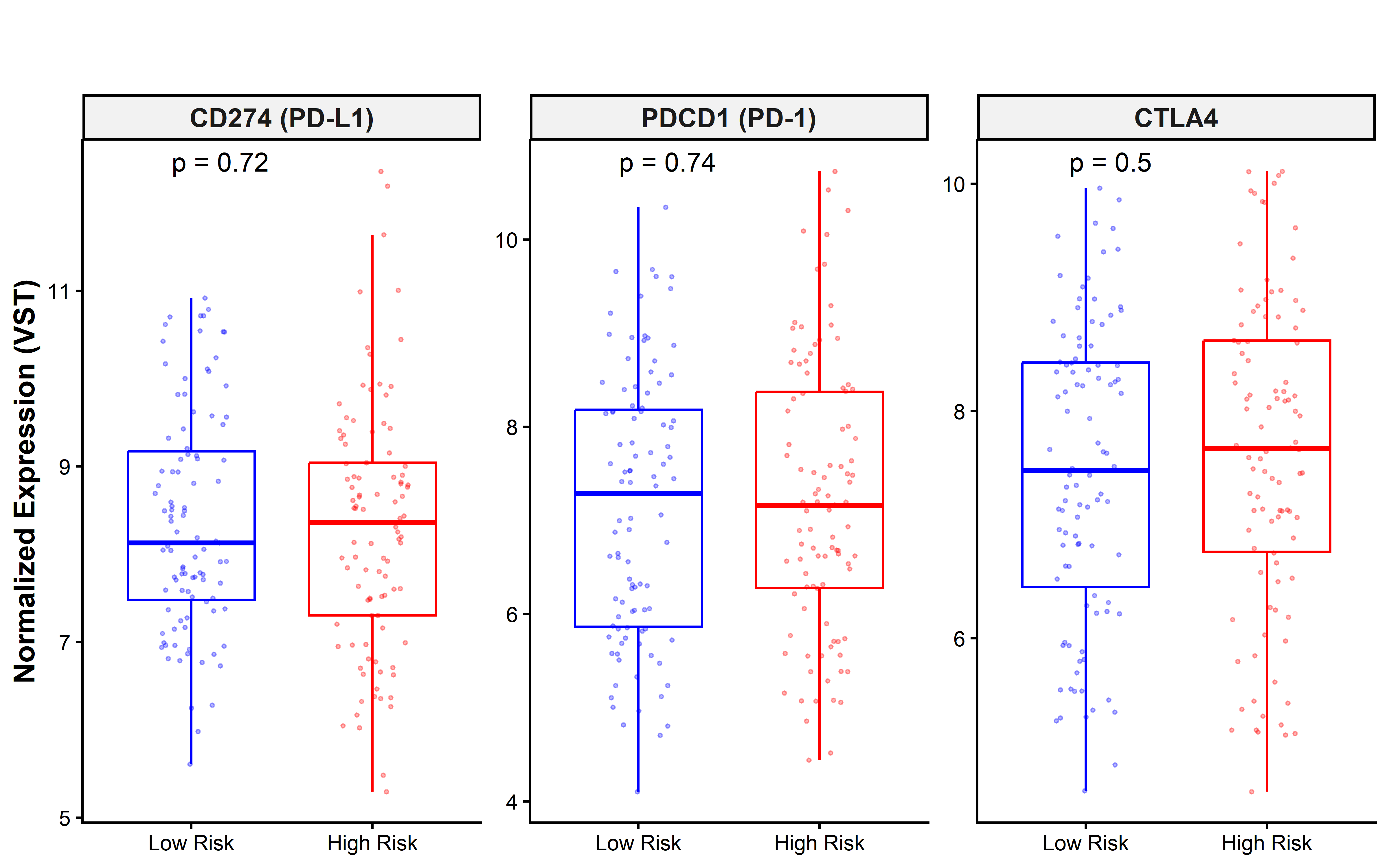
